# Vascularized Liver Tissue Embedded Bioprinting Utilizing GelMA/Nanoclay-based Composite hydrogels

**DOI:** 10.1101/2023.11.16.567470

**Authors:** Nima Tabatabaei Rezaei, Kartikeya Dixit, Hitendra Kumar, Jacob John, Giovanniantonio Natale, Simon S. Park, Keekyoung Kim

## Abstract

As the aging population grows, the need to regenerate non-self-repairing tissues becomes increasingly crucial for enhancing our quality of life. Tissue engineering offers a promising solution, particularly in recreating the intricate networks of blood vessels crucial for tissue vitality. These tissues rely on effective nutrient and oxygen circulation, with an optimal oxygen diffusion range of 100–200 µm. Yet, crafting vascularized in vitro tissues remains a significant challenge. This study addresses the challenge by using GelMA-based hydrogels as a photocrosslinkable support bath, a biocompatible and versatile choice for biological applications. To enhance the rheological properties for in vitro tissue engineering, Laponite (LPN) is introduced as a rheology modifier. The study optimizes the GelMA-LPN nanocomposite hydrogel composition, ensuring the desired physical, mechanical, and rheological properties, including recovery. The research also explores the biological implications, encapsulating liver cells within the nanocomposite hydrogel, and studying their behavior under perfusion conditions. This research presents a promising avenue for creating vascularized in vitro tissues, potentially advancing tissue engineering and regenerative medicine.

## 1. Introduction

Due to increasing aging population, the quality of our lives seems to rely on our capacity to regenerate impaired or ill tissues that lack the inherent ability to self-repair or renew into fully functional organs. As a result, the field of tissue engineering has provided possible solutions for this purpose [1,2]. Most of the body tissues are supplied with complex network of blood vessels, and their effective operation hinges on the circulation and diffusion of nutrients and oxygen, as well as the waste removal through nearby blood vessels. In fact, the optimal diffusion range for oxygen is typically cited as 100–200 µm, varying by tissue type. Beyond this threshold, the viability of cells seems to face challenges. For multicellular organisms to exceed this size, they need to stimulate the formation of new blood vessels through processes known as vasculogenesis and angiogenesis [3,4]. Following extensive years of research on engineering and regenerative medicine, a challenge is still remaining that is creating and constructing vascularized, sizeable in vitro tissues (>1 cm3) suitable for biological investigations and transplantation. Consequently, the need for bioengineered blood vessels remains on the rise, while the existing avenues for vascular design remain constrained [5].

Creating blood vessels within engineered tissue structures remains a challenge in the field of tissue engineering. In the past, when designing frameworks for tissue regrowth, the main focus was on making the materials, structure, and strength of these structures better to help cells grow [4,6]. In the past decades, researchers have tried out different methods to create artificial blood vessels for tissue engineering [7]. Some common ways to make these blood vessel-like structures include electrospinning [8], molding [3], rolling thin sheets [9], and using decellularization [10]. Advances in 3D printing methods have opened possibilities for creating blood vessel grafts containing cells and also complex vascularized tissue models [11,12]. So far, the technique of 3D bioprinting has been used to create two types of blood vessel structures: (a) vascularized tissue constructs, which are larger frameworks with incorporated channels or interconnected vessel systems, and (b) free-standing vascular tube-like structures [13]. Under the former category is indirect printing of perfusable matrices with sacrificial materials. In general, the process involves placing sacrificial materials within solid scaffolds, and these materials are later taken out to form empty pathways for adding cells. Typically, two different types of hydrogels are used. One acts as a support scaffold or sacrificial template, while the other serves as fugitive for creating the vasculature structure. When the sacrificial material is 3D printed as a network of blood vessels within a hydrogel matrix, changes in temperature or the use of specific solvents can dissolve the sacrificial template. This results in the creation of hollow vessel-like channels with specific entry and exit points for fluid flow. Fugitive inks are primarily chosen to have a reversible crosslinking mechanism. Examples of such materials include Pluronic F127, gelatin, agarose, and alginate [14–17]. Among these, Pluronic F127 hydrogel stands out due to its thermo-responsive nature. It maintains a stable viscosity over a wide sol–gel transition temperature range (10°C to 40°C), allowing it to remain consistent from room temperature to the body’s natural temperature [18,19]. However, by cooling it to 4°C, its elasticity decreases to liquifies and being able to be washed away easily. While there have been notable advancements in indirect bioprinting, creating extensive and intricate vascularized tissues remains challenging. This difficulty arises from the current hydrogel’s inadequate strength and rheological behaviors to serve as supporting bath, which hinders its ability to effectively uphold the entire structure [20]. On the other hand, to have a permanent supporting structure for the hollow networks, they should be properly crosslinked. Utilizing photocrosslinking is a resilient method to facilitate rapid, noninvasive, and precise gelation within 3D bioprinting which makes it easy to fabricate complex shapes and 3D structures out of low viscosity range biomaterials [21,22].

Photocrosslinking stands as a resilient strategy for providing rapid, noninvasive, and precise gelation with spatial and temporal control in the 3D bioprinting area [23,24]. The primary categories of photocrosslinking reactions employed in stereolithography (SLA) and digital light processing (DLP), as methods under 3D bioprinting, include the polymerization of (meth)acrylates, the thiol-yne (methacrylate) click reaction, and the dimerization of tyrosines [25]. Through various photocrosslinkable hydrogels, gelatin methacryloyl (GelMA), has gained considerable attention in tissue engineering and biomedical community due to its inherent controlable bioactivity and physicochemical [26,27]. Considering GelMA’s various biological attributes, it presents itself as a promising contender for deployment as a supportive bath material in embedded extrusion bioprinting (EEB). Nevertheless, the key concept in the EEB technique revolves around the selection of yield stress-fluids (YSFs), which are notably the preferred and widely adopted support bath materials due to their swift stress-triggered transition from liquid to solid state [28,29]. In order to make the GelMA suitable to be used as supporting bath, its rheological properties should be tuned. It was utilized in different applications of tissue engineering from composite hydrogel printing to support-bath material [30–32]. Laponite (LPN), a synthetic nanoclay with chemical formula of Si_8_Mg_5.45_Li_0.4_O_24_Na_0.7_. Structurally, it takes the form of a disc-like structure, 25 nm in diameter and 1 nm in thickness. Negative charges (OH−) are distributed on its surfaces, while positive charges (Na+) reside along its edges. Thanks to LPN’s biocompatibility, affordability, wide availability, thermal stability, processability, insensitivity to ions, and distinctive anisotropic behavior, LPN emerges as a promising modifier of rheology. Cai et al. [33] investigated the influence of adding LPN to alginate-gelatin composite hydrogel’s printability. Additionally, Munoz-Perez et al. [34] showed the impact of LPN on mechanical and rheological properties of Alginate. Also, LPN has been used as additive in gelatin or GelMA-based hydrogels. Barros et al. utilized GelMA-LPN for tumor modeling and studied the scaffolds rheological properties. It can also serve as a mechanical reinforcement component and crosslinker in various hydrogel systems [35,36].

In this study, we employed GelMA-based hydrogels as the photocrosslinkable support bath, serving as a matrix for liver tissue. However, GelMA alone lacked the desired rheological properties for our intended application. To address this, we utilized LPN as a rheology modifier to enhance the rheological characteristics and transform GelMA into a suitable material for embedded bioprinting purposes. Our research involved the optimization of the LPN content with a focus on achieving the desired physical, mechanical, and rheological properties, including recovery. Once the optimal composition was determined, we proceeded to assess its suitability for embedded printing of sacrificial material, the crosslinking process, and subsequent perfusion. Finally, in order to evaluate the biological aspects, liver cells were encapsulated within the nanocomposite hydrogel, and their behavior and fate were thoroughly examined under perfusion conditions.

## 2. Materials and Methods

### 2.1 Preparation of GelMA-LPN support bath

The synthesis of GelMA was conducted according to a previously reported protocol [37]. In a summary of the procedure, 5 g of powdered gelatin sourced from porcine skin (Type A, with a Bloom strength of 300, obtained from Sigma-Aldrich, St. Louis, MO, USA) was dissolved in 50 mL of RO purified water. Once complete dissolution was achieved at a temperature ranging between 48-50°C, 9-10 mL of glycidyl methacrylate (GMA) (Sigma-Aldrich) was added to the gelatin solution drop by drop. The solution was maintained at a temperature of 48-50°C with continuous stirring at 750 RPM for a duration of 12 hours.

Upon the completion of the reaction, the solution was transferred into dialysis tubing with a molecular weight cutoff of 12–14 kDa (provided by Fisher Scientific, Waltham, MA, USA) for dialysis against deionized water over the course of 3 days. The water was changed twice daily during this dialysis process. After the 3-day dialysis period, the remaining solution was frozen and subsequently subjected to lyophilization at -84°C to yield a foamy solid of GelMA. The lyophilized samples were stored at -20°C for future use.

To prepare the GelMA-LPN nanocomposite hydrogel, a stock solution of 10% (w/v) GelMA in PBS (referred to as stock 1) was prepared. Additionally, stock LPN solutions with concentrations of 0%, 1%, 2%, 3%, and 4% (w/v) in PBS were prepared, and vigorous agitation was employed for 3 hours to ensure the exfoliation of the nanoclay (this mixture is referred to as stock 2). Combinations as outlined in **Table 1** were prepared by mixing stock 1 and stock 2 in a 1:1 ratio and vortexing them thoroughly. Subsequently, the prepolymer solutions were incubated in a 37°C water bath overnight before being utilized for 3D printing applications.

**Table 1.**
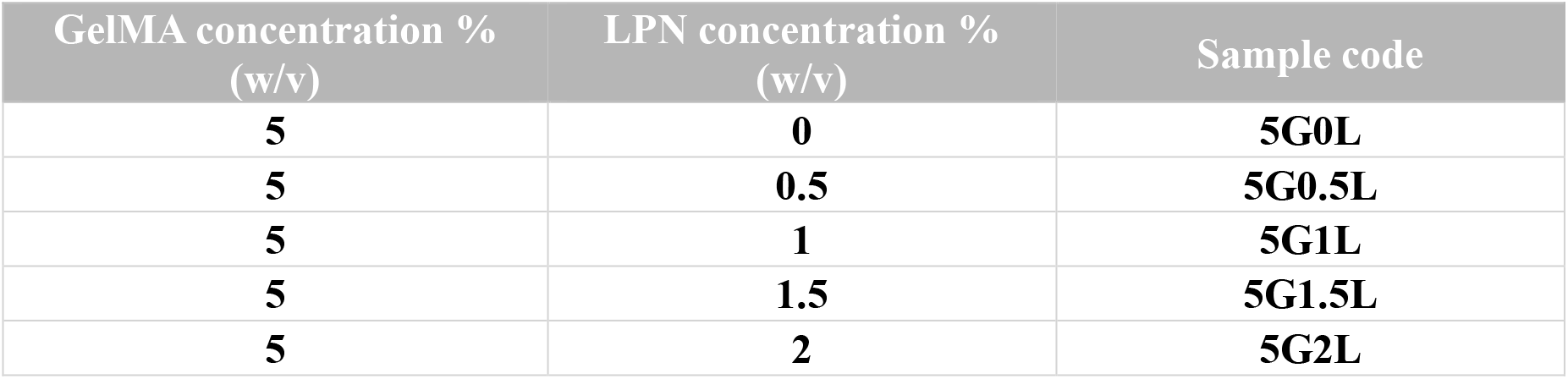
Different combinations used in this study to investigate the proper composition for supporting bath.

### 2.2 Mechanical, swelling and degradation measurements

The mechanical properties of the nanocomposite hydrogels were evaluated by measuring the compression modulus, which provides insight into the microstructural stiffness. To prepare the samples, varying concentrations of LPN were prepared and dispensed into molds (0.8 mm in diameter and 0.8 mm in height) with 500 μl of each sample added to individual wells. The prepolymer solution was subsequently crosslinked using 405 nm light for a duration of 1 minute to ensure uniform crosslinking throughout the entire section due to the large volume of the samples.

Subsequently, a vertical axis micromechanical testing machine, equipped with the custom-made flat-ended rigid cylinder with a diameter of 12.5 mm served as the probing instrument for compression testing, was utilized to record force-displacement plots for the hydrogels during indentation. Approximately 70% displacement was applied to the hydrogel surface with the speed of 20 mm/min to generate the force-displacement plot. Employing the initial dimensions and a in-house MATLAB code, the compression modulus was calculated based on the slope within the initial 10% strain region (linear region).

Measuring the swelling ratio of the crosslinked hydrogel samples is instrumental in assessing their water uptake capacity. In brief, 250 µl of the nanocomposite hydrogel solutions were cast into molds with a diameter of 0.8 mm. Subsequently, they were crosslinked under 405 nm light for a duration of 1 minutes. Following crosslinking, the hydrogels were immersed in PBS and placed in a 37°C incubator for 24 hours to allow for full hydration. Once completely hydrated, the weight of each sample in its hydrated state (w_w_) was recorded. Consequently, the hydrated hydrogel samples were frozen in a -80°C freezer and subjected to lyophilization for a period of two days to determine their dry weight (w_d_). Finally, the **equation (1)** was employed to calculate the swelling ratio, with five replicates.

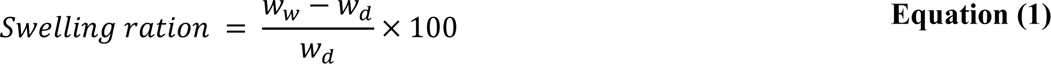

To investigate the degradation behavior of the bioinks, 750 μl of GelMA-LPN hydrogels were cast into 96-well plates and cured under 405 nm light for 1 minutes. All samples were subsequently lyophilized for 48 hours and weighed to obtain their initial weight (W_0_). Next, they were immersed in PBS at 37°C for 24 hours to achieve swelling.

The swollen samples were then placed in a 48-well plate containing a solution of collagenase I enzyme (50 U/mL) and incubated in a constant-temperature incubator at 37°C for degradation experiments. Samples (n = 3) were retrieved at 1, 2, 3, 4, 6, 9, 12, 24, and 48 hr. Subsequently, each sample was subjected to 24 hours of lyophilization to determine the remaining weight (W_r_). The biodegradation property (Q_d_) of the hydrogels was assessed as the percentage of the remaining weight of the disks after degradation, using **equation (2)**.

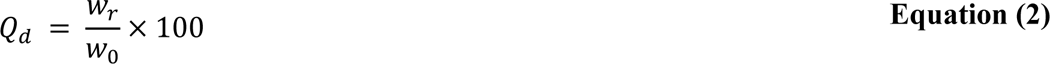

### 2.3 Fourier Transform Infrared (FTIR) Spectroscopy

The different combinations mentioned in section 2.1 were used after incubation at 37 °C followed by freezing at -80 °C and lyophilization. FTIR spectra were determined with a Thermo Nicolet Avatar 370 Fourier Transform Infrared spectrometer. Each sample was subjected to 32 scans per sample at 1.9 cm^−1^ resolution from 4000 to 475 cm^−1^.

### 2.4 Scanning Electron Microscopy (SEM) and porosity quantification

To examine the microstructure and pore size of the GelMA-LPN hydrogel, scanning electron microscopy (SEM) was employed. For this characterization, 250 µl of hydrogel precursor was prepared for each hydrogel group with specific concentrations and crosslinked using 405 nm light in cylindrical container. Following crosslinking, the samples were frozen in a -80°C freezer and subjected to a two-day lyophilization process. Once lyophilized, the samples were sputter-coated with gold using a sputtering machine and subsequently imaged using a SEM (phenom ProX).

### 2.5 Rheological Characterization

All rheological characterizations were conducted using an MCR 302e rheometer (Anton Paar, Austria) equipped with a Peltier plate for temperature control. A stainless-steel parallel plate with a diameter of 25 mm and a gap distance of 0.5 mm at room temperature was employed for all experiments. Various rheological tests were performed to investigate the flow and viscoelastic behavior of the support-baths. Gel yield stresses and the linear viscoelastic (LVE) regions were determined by conducting strain sweeps ranging from 0.01% to 100% at a constant angular frequency of 10 rad/s. Oscillatory angular frequency sweeps were carried out in the range of 0.1–100 rad/s at a constant 10% strain to monitor the dynamic rheological behavior. Shear rate sweeps were conducted to monitor the shear-thinning behavior and viscosity alterations of the formulations over a range of 0.01–100 1/s. To investigate the recovery behavior of the support-bath during hydrogel extrusion, cyclic strain tests were performed at low and high oscillatory strains of 10% and 500%, respectively, at a constant angular frequency of 10 rad/s, with each cycle lasting 10 seconds. To record the photocrosslinking kinetics, a transparent glass plate was integrated into the setup, allowing for the illumination of the hydrogel with 405 nm light from the bottom of the parallel 25 mm geometry. The light source, was initiated after 1 min of the beginning of the experiment and continued for a duration of 1 minute. Oscillatory measurements were conducted with a 1% shear strain and a frequency of 1 Hz, and both the storage and loss moduli for the photocrosslinking bioink were simultaneously recorded.

To explore the time-dependent rheological characteristics of both the 5G0L and 5G1.5L samples, an assessment of gel yield stresses and LVE regions was conducted. This evaluation involved conducting strain sweeps over a range from 0.01% to 100% at a consistent angular frequency of 10 rad/s. The measurements were performed using a 25 mm geometry and a 0.5 mm gap. Two distinct conditions were considered for each sample: one taken immediately after vortexing and another after allowing the samples to rest for 15 minutes. The objective was to compare and analyze the material’s behavior under these two conditions, shedding light on the time-dependent rheological properties.

### 2.6 Supporting bath transparency evaluation and SLA printing

Transparency assessments of the hydrogels were carried out employing UV–Vis spectroscopy (Shimadzu 2600). Square-bottom cuvettes with a width of 5 mm were employed for the measurements, covering the wavelength range from 350 to 950 nm. Before conducting the measurements, the samples underwent a preparation process. They were vortexed and subsequently centrifuged at 1500 rpm for 1 minute to eliminate any air bubbles and ensure uniformity in the samples for accurate transparency measurements.

Following the preparation of the GelMA-LPN solution as outlined in section 2.1, the intended structures were fabricated by subjecting the prepolymer solutions to near UV light (wintech PRO4500) (with a wavelength of 405 nm and an intensity of 128 mW cm−2). This was achieved using a customized SLA bioprinting system, as previously described in earlier reports [38,39]. The exposure duration for each bioink combination ranged from 5 to 7 seconds. To enable the crosslinking of the entire cross-section simultaneously and create the designed structures, circular molds with a depth of 450 μm were employed. After the crosslinking of the structures, any remaining uncrosslinked portions were thoroughly washed with PBS (Phosphate-Buffered Saline). Subsequently, the printed structures were colored with pink food dye to enhance their visibility for photography and visual documentation.

### 2.7 Embedded extrusion printing evaluation

In this study, we employed an in-house modified Fused Deposition Modeling (FDM) printer, which was equipped with an air dispenser for ink extrusion. Our printing process utilized a nozzle with a diameter of 200 µm. Pluronic F-127 was chosen as the sacrificial material. PF-127 is a thermoresponsive polymer, exhibiting a fluid-like behavior at lower temperatures and transitioning to a gel-like state at temperatures above room temperature (as depicted in **Figure S6**). After experimentation with different compositions, a 30% (w/v) PF-127 solution was determined to be the most suitable concentration of the sacrificial material. To enhance visibility, we added Eosin-Y dye to the PF-127 solution. To assess the printing of the sacrificial material within the bath material, a custom G code was developed. This G code instructed the printer to create five straight lines, each measuring 30 mm in length. The printing speeds for these lines ranged from 200 mm/min to 600 mm/min, with 100 mm/min intervals (please refer to the supplementary videos for more details). Furthermore, the extrusion pressure was varied from 20 psi to 60 psi, with 5 psi increments, while printing these five lines.

Further, a total of five structures were printed with different geometrical intricacies in the chips used as mold that were additively manufactured using clear resin (**Figure S7(A) and (B)**). After printing these structures, the bath material having the photoinitiator with embedded structures was photo crosslinked using a near UV light projector with specifications mentioned before for 15-20 seconds regarding the structures. Next, the chip was transferred to a refrigerator having ∼4-8°C temperature to liquefy the Pluronic F-127 channel for ∼1 hour. Later, a syringe pump was used to flush out the liquid PF-127 channel by perfusing cold PBS solution. After evacuating the Pluronic F-127 from the channels, it was perfused with media for more than 10 hours to check its perfusion capabilities (**Figure S7(C)**).

### 2.8 Evaluation of cell fate and proliferation in GelMA-LPN

The cells used in all experiments were HepG2 cell lines, cultured at 37°C in a 5% CO2 atmosphere. The cell growth medium consisted of Dulbecco’s Modified Eagle Medium (DMEM) (Lonza, Basel, Switzerland) supplemented with 10% v/v heat-inactivated fetal bovine serum (FBS) (Thermo Fisher Scientific, Waltham, MA) and 1% v/v penicillin-streptomycin (Sigma-Aldrich). The cells were cultured in T75 cell culture flasks (VWR International) and were only utilized or subcultured when they reached 85%–90% confluence.

To characterize and visualize cell proliferation within GelMA-only and GelMA-LPN bioinks, cells were encapsulated within hydrogel scaffolds and cultured for a duration of one month. A hydrogel prepolymer solution comprising 5% w/v GelMA, which served as the control sample, and 5G1.5L (identified as the optimal concentration for use as a support matrix) was prepared containing LAP in PBS. Cells were added at a density of 2.6 × 10^6^ cells ml^−1^ to the prepolymer solution and uniformly dispersed. The final bioink was pipetted into a mold with a depth of 450 μm, using approximately 1000 µl of the solution.

Subsequently, the cell-laden prepolymer solution was crosslinked by exposing it to 405 nm light in the previously mentioned SLA bioprinting system for 15 seconds to ensure complete crosslinking. Following crosslinking, the sheets were transferred from the mold to a Petri dish, washed with PBS, and immersed in 5 ml of fresh growth media. The samples were then incubated at 37°C under a 5% CO2 atmosphere. To investigate cell adhesion and morphology, the encapsulated samples were punched using a 5 mm surgical punch, and at least three disks from each sample were stained with phalloidin (Cytoskeleton, Denver, CO, USA) and 4’,6-diamidino-2-phenylindole (DAPI) (FluoroshieldTM with DAPI, Sigma-Aldrich) for cytoskeleton and nuclei staining, respectively, at time points of 7, 10, 14, 21, and 28 days of culture. The staining procedure involved washing the hydrogel samples with PBS three times before transferring them to a 24-well plate. They were then fixed in 4% v/v paraformaldehyde for 90 minutes. After fixation, the samples were washed with PBS and treated with 0.5% v/v Triton X-100 in PBS (Sigma-Aldrich) for 20 minutes to permeabilize the cell membranes. Subsequently, 500 µl of phalloidin 488 stock solution was added to the samples and incubated at room temperature for 90 minutes. Afterward, the samples were washed three times with PBS. The samples were then transferred to a glass slide, and DAPI stock solution was applied. A cover slip was placed on top, and the samples were imaged using 10× and 20× objective lenses in an inverted fluorescence microscope (ECHO revolve) equipped with DAPI and EGFP channels.

A minimum of five images per sample were analyzed using ImageJ software. This analysis aimed to precisely quantify both the size of the clusters and the number of clusters within a given unit of area.

### 2.9 Embedded in-gel bioprinting evaluation

Human hepatocellular carcinoma (HepG2 cells obtained from ATCC) were cultured in Dulbecco’s modified Eagle medium (DMEM) supplemented with 10% fetal bovine serum (FBS) and 1% penicillin/streptomycin (PEST). Adherent cells were trypsinized, centrifuged, and resuspended in 5G1.5L hydrogel at a concentration of 2.5 × 10^6^ viable cells ml^−1^.

The cell-laden hydrogel was then introduced into the previously sterilized mold, and a structure similar to the one depicted in **Figure 7(A)** was bioprinted using sterilized PF127. After the crosslinking process, the bioprinted structure was stored in 4-8°C for 30 minutes to allow the PF127 to liquefy. Subsequently, by perfusing the culture media, the PF127 was washed out. After immersing the bioprinted construct in the culture media, the entire perfusion system was incubated at 37°C in a 5% CO2 environment for a week.

Multiple chips were manufactured to enable sample collection at both day 3 and day 7 following perfusion. Slices, with a thickness of approximately 1.5 mm, were taken from both the section containing the hollow channel and the solid section to facilitate a comparison of cell viability. These cross-sectioned samples were washed twice with PBS to remove the culture media before being incubated in a live/dead assay staining solution (obtained from Biotium, Hayward, CA, USA) for a duration of 30 minutes. The assayed samples were imaged using a fluorescence microscope (ECHO revolve), and efforts were made to keep the samples hydrated to maintain consistency in the measurements.

### 2.10 Statistical analysis

All data are presented as mean values accompanied by the standard deviation. Statistical analysis was conducted utilizing Minitab. To determine statistical significance, a one-way analysis of variance was employed along with Tukey’s multiple comparison test. Differences were regarded as statistically significant when the p-value was less than 0.05, and the significance levels were indicated as follows: ns (not significant) for p > 0.05, *p < 0.05, **p < 0.01, ***p < 0.001, and ****p < 0.0001.

## 3. Results

### 3.1. GelMA-LPN nanocomposite hydrogel preparation

To comprehend the interaction between GelMA and LPN, an examination of the chemical properties of these components was conducted. In its dry state, LPN exists in the form of microscale tactoids, characterized by dimensions ranging from 10 to 100 μm. These tactoids represent stacked layers of silicate, mediated by Na^+^ ions (**Figure S1(A)**) [40]. The negative charge on the surfaces of LPN is attributed to the partial substitution of Mg^2+^ ions with Li^+^ ions [41]. When LPN powder is dispersed in water, the tactoids undergo hydration, leading to the dissociation of Na^+^ ions. This results in a permanent negative charge on the particle surfaces and causes the exfoliation of the stacked structures into individual particles, as outlined in **Figure S1(A)**. The exfoliation process of LPN is primarily governed by the adsorption/desorption of Na^+^ ions from the tactoids during hydration, which can be influenced by the addition of salts to the solutions, such as in the case of PBS. When LPN nanodiscs are not fully exfoliated due to the presence of ions, when they are added to PBS, they tend to form clusters of stacked discs with dimensions exceeding 100 nm (**Figure 1(A)**) [42]. The addition of LPN in PBS before and after exfoliation is shown in figure S1a and b. It is evident that samples containing more than 1% LPN form a gel after 3 hours of exfoliation. The formation of stable gels depends on solid concentration and incubation time. LPN at concentrations more than 1% (w/v) form a physical gel within several hours due to the self-assembly of exfoliated nanodiscs into a house-of-cards structure through face-to-edge attractions, as presented in **Figure S1(A)** [43]. At lower nanoclay concentrations (≤1% w/v), the gel formation process is considerably slower. Previous studies have reported that LPN at concentrations less than 1% (w/v) may form a gel after several months [44].

**Figure 1.**
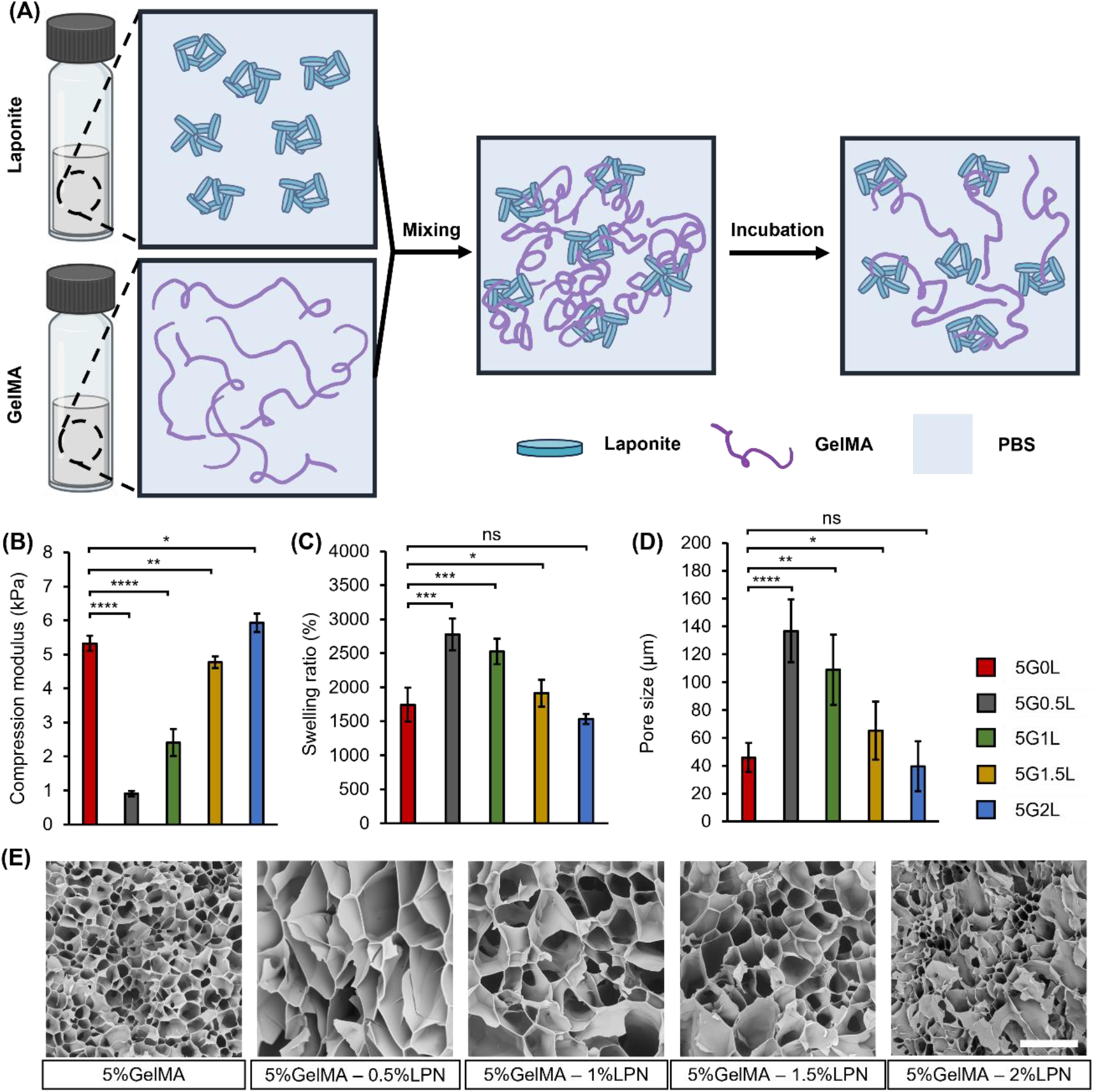
Physiochemical and mechanical properties evaluations of the GelMA-LPN nanocomposite hydrogels. (A) schematic of preparation of the GelMA-LPN hydrogel, (B) Compression modulus investigation of the crosslinked hydrogel samples, (C) Swelling evaluation of the hydrogels, (D) pore size measurement from SEM results representing microstructure of the hydrogels, (E) microstructural evaluation of the hydrogels samples (scale bar=200um)

When 5% GelMA is added to the exfoliated LPN solutions, **Figure S1(B)-iii** indicates that there are interactions between GelMA and LPN. **Figure 1(A)** illustrates the mechanism for the synthesis of nanocomposite hydrogels containing LPN and GelMA, as previously reported by Li et al. [45]. In the initial step, GelMA solution was added to a dispersion of LPN. It is noticeable that immediately after mixing, the solution transitions from transparency to opaqueness (**Figure S1(B)-iii**). This change can be attributed to the negatively charged surfaces of LPN particles and the strong polyampholytic properties of GelMA, which make it easily polarized by a charged surface. Such polarization can induce attractive electrostatic interactions between GelMA and LPN. Additionally, these mixtures are in a semi-stable state, and the strong interaction between the components can lead to spinodal decomposition, resulting in a less transparent solution [46]. However, in the subsequent step, when the mixture is left to incubate in a water bath at 37 °C overnight, the turbidity of the mixture significantly decreases (**Figure S1(B)-iv**). This reduction is likely due to the weakening of the non-covalent interactions between the nanoclay and gelatin, preventing the precipitation of the mixture when heated to 37 °C.

FTIR spectra were utilized to detect the functional groups and structural changes in the GelMA-LPN nanocomposite hydrogels at a molecular level. **Figure S1(C)** displays the FTIR absorption bands of the GelMA-LPN nanocomposite hydrogels. The two characteristic peaks at 1645 and 2935 cm^−1^ correspond to the C═O stretching in the carboxamide functional group and C—H stretching, respectively. The intense peak at 999 cm^−1^ is associated with Si—O stretching in the nanoclay, and the absorption band around 3317 cm^−1^ can be attributed to the —NH stretching in the amide group and —OH stretching in Laponite [45,47]. Furthermore, no new vibrations were observed, indicating the absence of new covalent bonds between the clay and the polymers in the nanocomposites after the addition of gelatin.

### 3.2. Mechanical and physiochemical properties investigation of GelMA-LPN nanocomposite hydrogel

Due to influence of the scaffold’s mechanical properties on the cell proliferation and fate [48,49], it is one of the necessary steps to evaluate the final structures stiffness. **Figure 1(B)** is showing the influence of adding LPN to GelMA hydrogel. The compression modulus of the 5% GelMA was 5.32 ± 0.22 kPa and by adding 0.5% LPN, the compression modulus of the hydrogel changed to 0.9 ± 0.08 kPa (decreased by 5.8 folds). Future increasing the LPN concentration resulted in increasing the compression modulus up to the relatively same level as 5G for the sample containing 1.5% and 2% LPN which is around 5kPa and is proper to be used as 3D scaffold model.

In addition to the mechanical properties, swelling characteristics would have great impact on the solute diffusion and nutrition circulation within the scaffold [50]. **Figure 1(C)** is reporting the water absorption capability of the crosslinked Hydrogels with different combination of GelMA and LPN. It is reported that the addition of the LPN will first increase the swelling ratio from 1743.3 ± 247.59% to 2776.69 ± 233.76% for 5G and 5G0.5L samples respectively. However, for the samples with LPN concentrations more than 0.5%, the swelling ration keeps decreasing to 1535.68 ± 75.1% for 5G2L hydrogel sample.

Both the mechanical and swelling properties are representing structural characteristics. In order to prove the reported properties, internal 3D structure of crosslinked scaffolds was visualized through SEM. **Figure 1(D)** and **(E)** are representing quantified pore size and SEM images respectively. 5G sample is showing comparatively small and uniform pore size of 45.79 ± 10.44 μm which was increased to 136.73 ± 22.46 μm for 5G0.5L sample as the one having lowest compression modulus and highest swelling ratio that are in correlation with each other.

Biodegradability stands as a highly desirable attribute within the tissue engineering area, particularly concerning hydrogel materials. It serves as an indicator of the scaffold’s ability to maintain its structural integrity over time. Given the inherent slow degradation of hydrogels in a PBS environment, collagenase enzyme has been employed to expedite the degradation process and explore the degradation profiles of the various samples. As depicted in **Figure S2**, the 5G sample exhibits complete degradation within a brief span of 5 hours. Notably, the sample enriched with 0.5% LPN demonstrates the most rapid degradation rate in comparison to the other formulations. In stark contrast, samples incorporating 1.5% and 2% LPN within their composition manage to retain 55.69% and 60.93% of their initial dry weight after 48 hours, respectively. This compelling outcome positions them as promising candidates for in vitro modeling applications, where extended structural stability is a valuable attribute.

### 3.3. GelMA-LPN rheological properties assessment and optimization as supporting bath

The successful implementation of suspended printing is relied upon the availability of a material with optimized rheological properties. Specifically, the bath material should not only exhibit Bingham behavior, which facilitates the printing process, but also demonstrate a stable sol-gel transition to maintain the integrity of the constructed structures [51,52]. In order to assess the suitability of GelMA-LPN hydrogel as a supporting bath material, we conducted rheological assessments on various GelMA and LPN combinations. These hydrogels were subjected to a range of amplitudes of oscillatory shear stress, allowing us to measure both the elastic storage modulus (G’) and the loss modulus (G’’).

The dynamic moduli of 5% GelMA composites, containing 0%, 0.5%, 1%, 1.5%, and 2% LPN, are presented during the strain amplitude (**Figure 2(A)**) and angular frequency sweeps (**Figure 2(B)**). As the LPN concentration increased, G’ and G’’ in the linear viscoelastic regime increased. Figure 2(A) reveals the yield strain of the materials, where a transition occurs from an elastic gel state (G’ > G’’) to a viscous liquid-like state (G’’ > G’) with increase in shear strain [53,54]. It has been observed that the control and the sample containing 0.5% LPN exhibit no crossover between G’ and G’’. However, at higher LPN concentrations, they demonstrate distinguishable yielding points occurring at lower shear strains as the LPN percentage increases.

**Figure 2.**
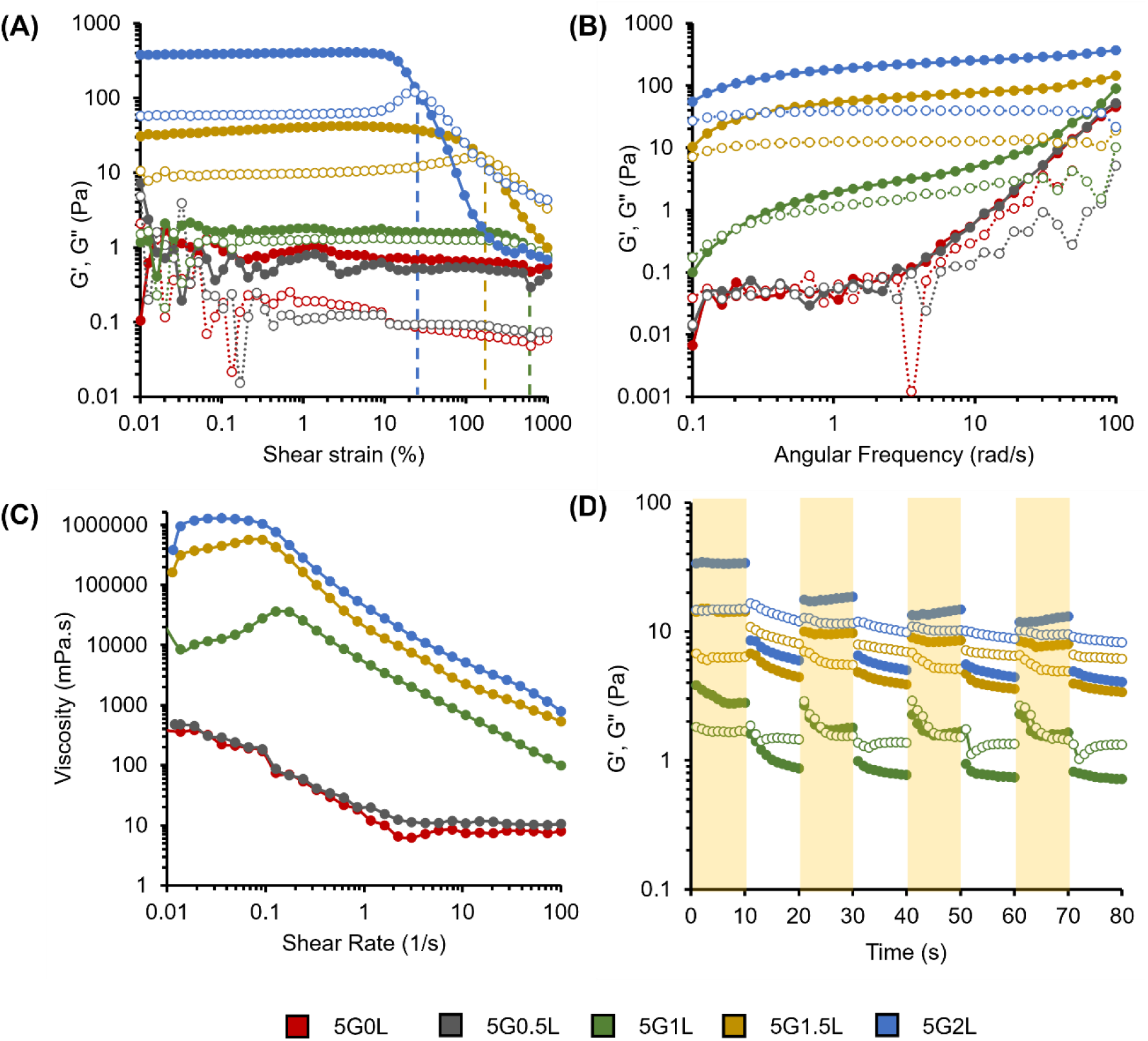
Dynamic rheological characterization of the support-bath representing the effect of LPN concentration on flow behavior and recoverability of the structure of nanocomposite hydrogels at room temperature (A) Strain amplitude sweep profiles of GelMA-LPN hydrogels, (B) frequency sweep profiles, (C) viscosity vs. shear rate plots revealing the shear thinning behavior of the support material, (D) cyclic strain and recovery measurements at high (500%) and low (10%) strains showing storage (G′) moduli of the samples in 4 cycles. Storage modulus (G’) (filled symbols) and loss modulus (G”) (open symbols).

The results from oscillatory shear tests indicate that composites with 1.5% LPN and higher display solid-like behavior (G’ > G’’) during the oscillation frequency sweep (**Figure 2(B)**). Conversely, the sample with 1% LPN exhibits liquid-like behavior (G’’ > G’) at frequencies below approximately 0.2 rad/s, while the samples with 0.5% and 0% LPN concentrations do not display a clear trend as observed in others as shown in **Figure 2(B)**.

Furthermore, the shear-thinning behavior of the composite support-bath materials was investigated through a shear rate sweep test (**Figure 2(C)**). As depicted in the graph, all tested concentrations exhibit a similar trend of viscosity reduction. However, with an increase in LPN concentration, the viscosity of the samples increases.

As a vital rheological property of the supporting bath, owing to its thixotropic characteristics, the disturbed matrix gradually rebuilds interactions, forming a network over time [54,55]. This self-recovery property is a crucial feature for a composite matrix intended for use as a support-bath material. **Figure 2(D)** illustrates the recoverability of the composite during cyclic deformation. As shown in the graph, even within a short period of 10 seconds, structures containing 1% and 1.5% LPN nearly return to their initial storage moduli. However, the composite with 2% LPN exhibits lower recovery in storage modulus compared to the starting point.

### 3.4. Transparency evaluation of the supporting bath

The optical transparency of hydrogels holds significant importance, particularly in specific biomedical applications where observation of conditions inside a hydrogel is essential [56]. Additionally, in the context of this study, it is crucial to have a support bath that is transparent. This transparency is imperative for visualizing the embedded printed structures and optimizing printing parameters. Furthermore, there are several studies that demonstrate the influence of hydrogel transparency on the resolution of light-induced bioprinting techniques [57,58]. Therefore, the evaluation of optical transparency becomes a pertinent characteristic to assess for the reported bioink in this study. As illustrated in **Figure 3(A)** and **(B)**, the 5% GelMA control hydrogel solution, devoid of LPN, exhibited remarkable transparency. Conversely, the introduction of LPN led to a gradual reduction in the solution’s transmittance.

**Figure 3.**
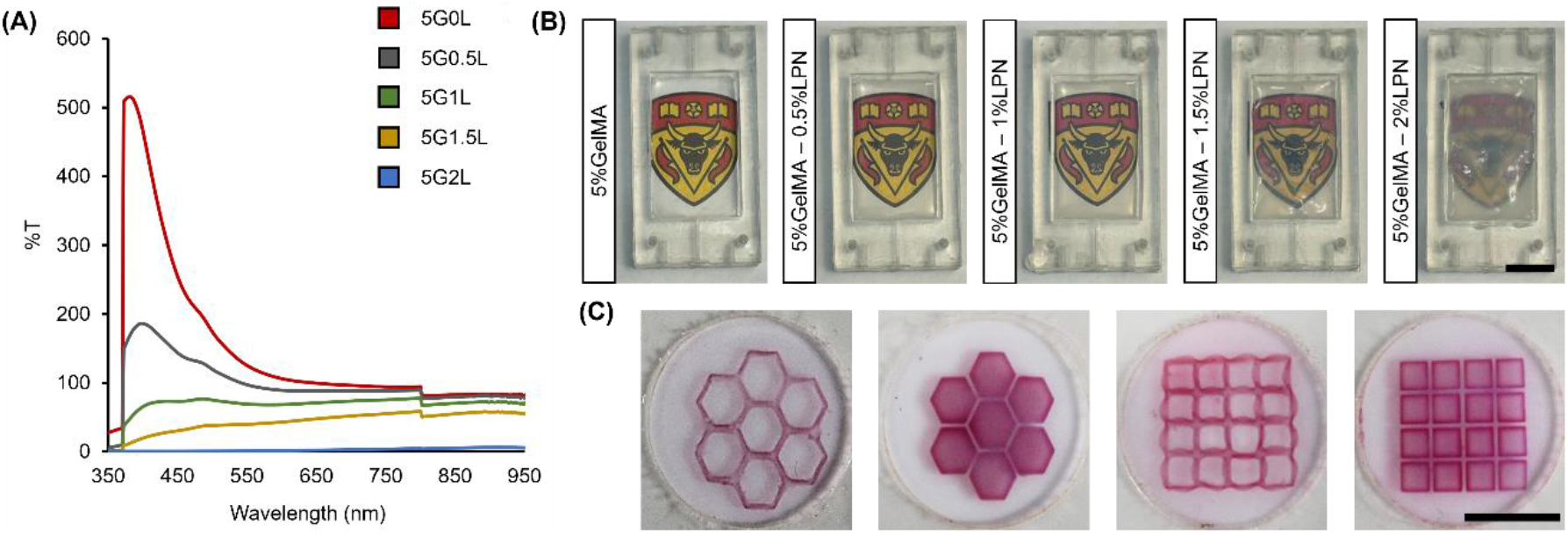
Transparency evaluation and SLA printing. A) transmittance of the support bath hydrogel in the UV and visible light range, B) Photos showing the transparency of the support bath in the mold being used for chip fabrication (scale bar: 1cm), C) Different structures crosslinked and colored to visualize the precision of SLA printing, using the GelMA-LPN support matrix hydrogel (scale bar: 1cm)

Moreover, in the context of employing GelMA-LPN nanocomposite hydrogels for photocrosslinking applications, diverse patterns were fabricated using UV light crosslinking to evaluate the precision and resolution of stereolithography-based patterning, as demonstrated in **Figure 3(C)**. These findings affirm the suitability of the provided bioink for use as a high-resolution photocrosslinkable bioink.

### 3.4. Freeform embedded printing evaluation within GelMA-LPN supporting bath

**Figure 4** illustrates the behavior of the sacrificial material within the GelMA-LPN supporting bath during the printing process. The graph depicts how variations in extrusion pressure and printing speed impact the creation of lines within the GelMA-LPN bath material. Initially, an extrusion pressure of 20 psi was found to be insufficient for the extrusion of the 30% (w/v) PF-127 due to its high viscosity at room temperature, as determined by viscosity testing. When the extrusion pressure was increased to 25 psi, it exceeded the yield stress required for the extrusion of PF-127, causing it to flow through the nozzle. The results also reveals that even at higher printing speeds, the extrusion of PF-127 did not consistently yield the desired line structure. At 25 psi, the only successful print occurred at a printing speed of 200 mm/min. Upon further increasing the extrusion pressure to 30 psi, over-extrusion occurred at lower printing speeds (200 and 300 mm/min), while successful prints were achieved at higher printing speeds (400, 500, and 600 mm/min). Similarly, at 40 psi, over-extrusion was observed at lower printing speeds, where the material extruded exceeded the amount required for line fabrication. This behavior was consistent at higher extrusion pressures, such as 50 and 60 psi.

**Figure 4.**
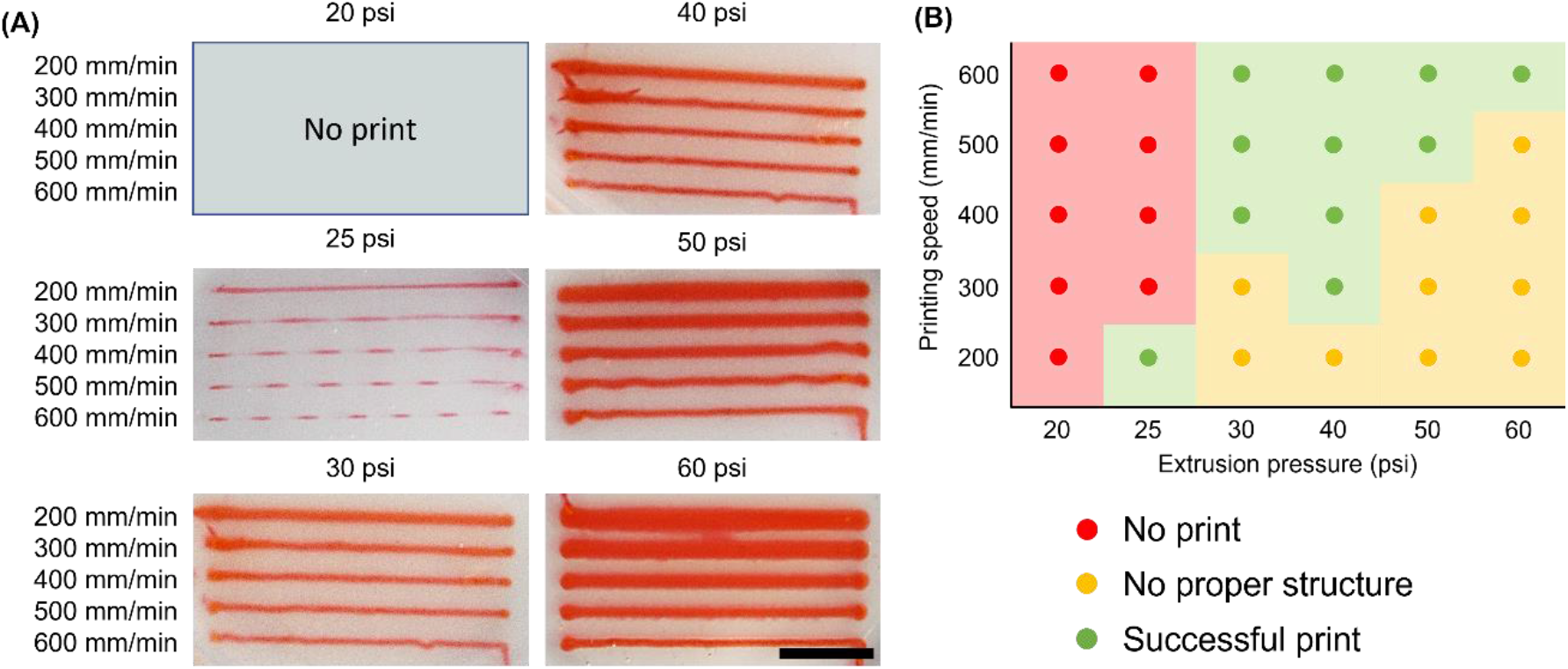
Printing conditions evaluation and optimization (scale: 10mm)

However, from **Figure 4**, it is evident that, at a given pressure, varying the printing speed can optimize the structure’s diameter to some extent. To obtain a structure with a diameter similar to the nozzle size, a trade-off between extrusion pressure and printing speed is necessary. Nevertheless, with hydrogels, achieving a diameter closely matching the nozzle size is challenging due to the die swelling effect or Barus effect, as discussed by Paxton et. al [59]. In sum, we utilized 25 psi extrusion pressure and a printing speed of 200 mm/min to print five 2D structures. These structures were later used to create hollow channels and perfused with media, as outlined in section 2.6. Printing videos of the structures shown in **Figure 4**, are presented in **supplementary videos 6-10**.

Under the optimized printing conditions, the efficacy of the newly developed support bath for embedded printing applications was assessed. A range of structures was successfully printed within the 5G1.5L support bath, followed by the crucial steps of crosslinking and subsequent perfusion. This process resulted in the creation of intricate hollow channels on the chip, as visually demonstrated in **Figure 5**.

**Figure 5.**
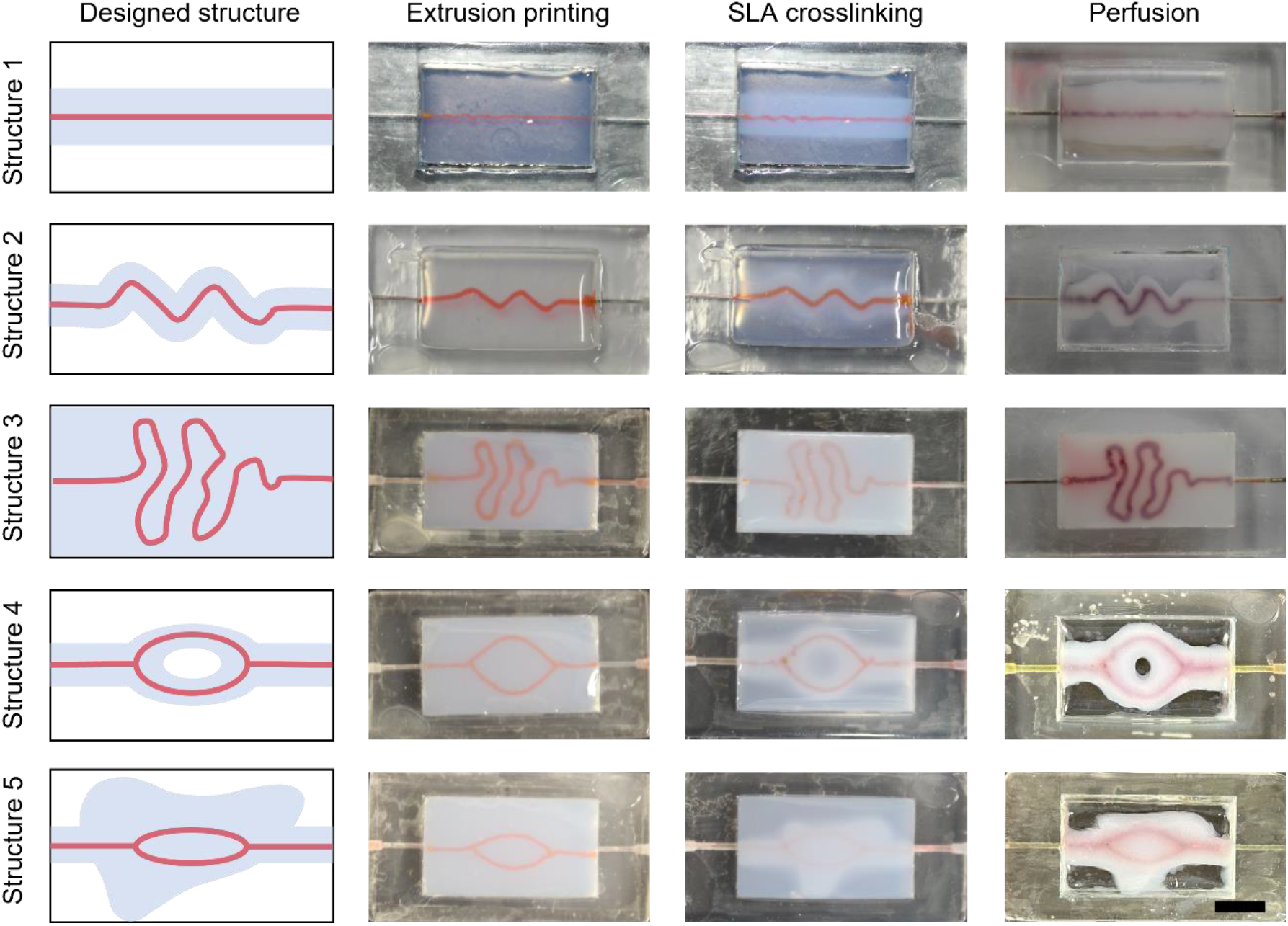
Vasculature printing and perfusion. Different structures being used for printing fugitive ink embedded in the GelMA-LPN support bath and post processing of photocrosslinking, washing the sacrificial ink and perfusion with colored distilled water (scale: 250 um)

### 3.5. Cell proliferation within bioprinted constructs

To assess the GelMA-LPN’s capability to maintain encapsulated hepatocytes with high viability and sustained functionality, we initially evaluated the bioink using HepG2 cells encapsulated within the hydrogel. In order to visualize the morphology and organization of cells within the GelMA-LPN hydrogel scaffolds, actin cytoskeleton and nuclei were stained at various time points during culture (**Figure 6**).

**Figure 6.**
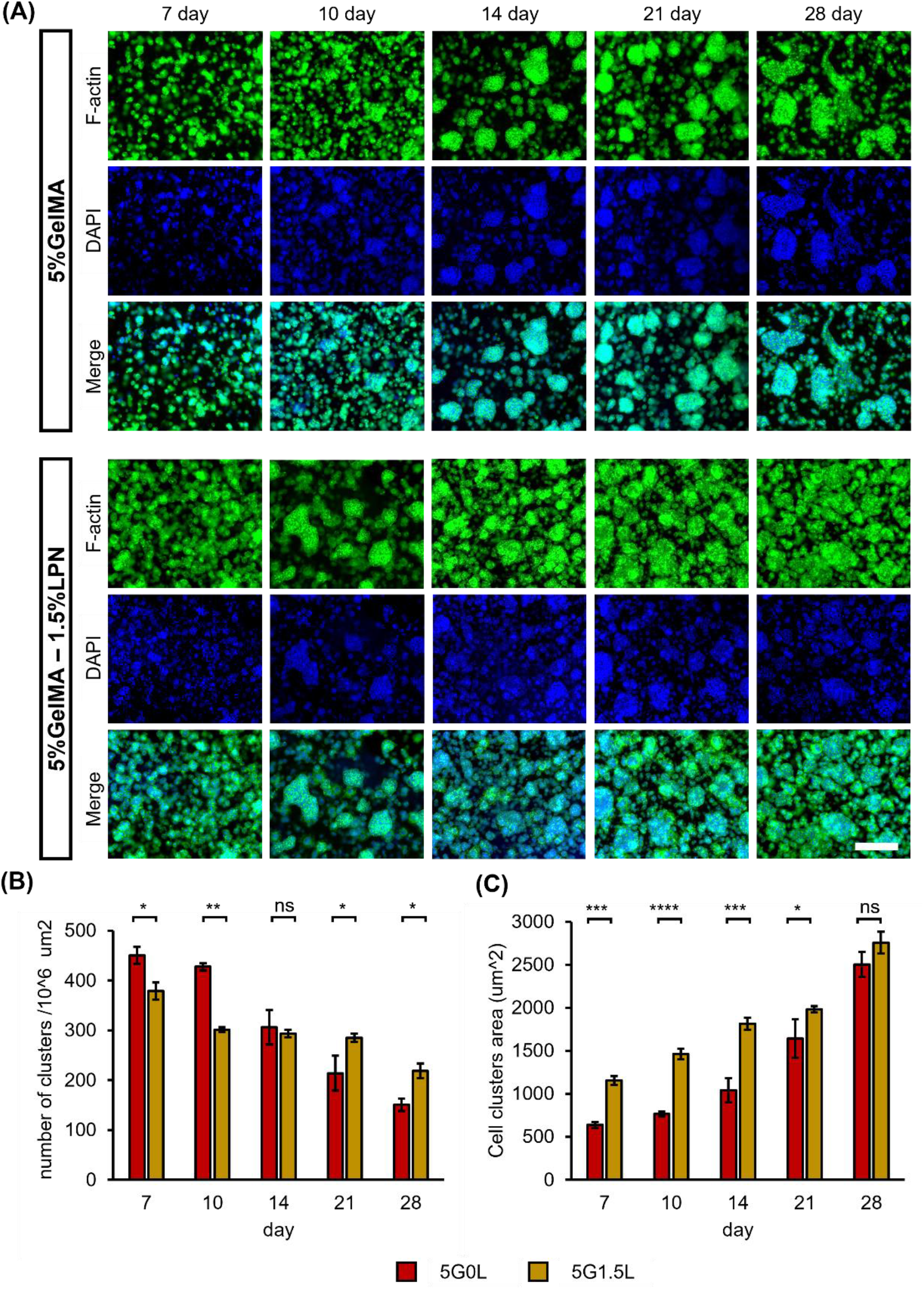
HepG2 cells proliferation evaluation within the GelMA-LPN hydrogels. A) Phalloidin-DAPI staining to visualize the cytoskeleton and cell nuclei to evaluate the proliferation of the cells in the period of one month for 5G0L and 5G1.5L samples (scale: 1cm)

Upon comparing cellular morphology between the control and the 5G1.5LPN sample, it became evident that the presence of LPN in the hydrogel led to more pronounced cluster formation at all observed time intervals, as compared to the control lacking LPN. In general, the clusters formed in the presence of LPN exhibited a more distinct and organized architecture of the actin filament-based cellular network. Moreover, they displayed multicellular aggregates that both increased in size and number throughout the culture period, spanning from day 7 to day 28. This trend was notably more pronounced in the hydrogel containing LPN (**Figure 6(A))**.

**Figure 6(B) and (C)** illustrate the number of clusters per unit area and the cluster area, respectively. These measurements further corroborate the increased formation of aggregates and the rapid expansion in cluster size within the 5G1.5LPN sample.

### 3.5. Cell proliferation within bioprinted constructs

Bioprinted constructs were cultured for one week under perfusion conditions. Viability assessments were conducted on both the samples from the channel-printed portion and the solid section. As depicted in Figure 7, it is evident that the cells along the channel wall exhibit a viability exceeding 95%. This can be attributed to the continuous perfusion of fresh media, providing an ample supply of nutrients and oxygen. In contrast, the viability in the solid section declined to less than 50%, primarily due to the limited availability of essential nutrition and oxygen.

**Figure 7.**
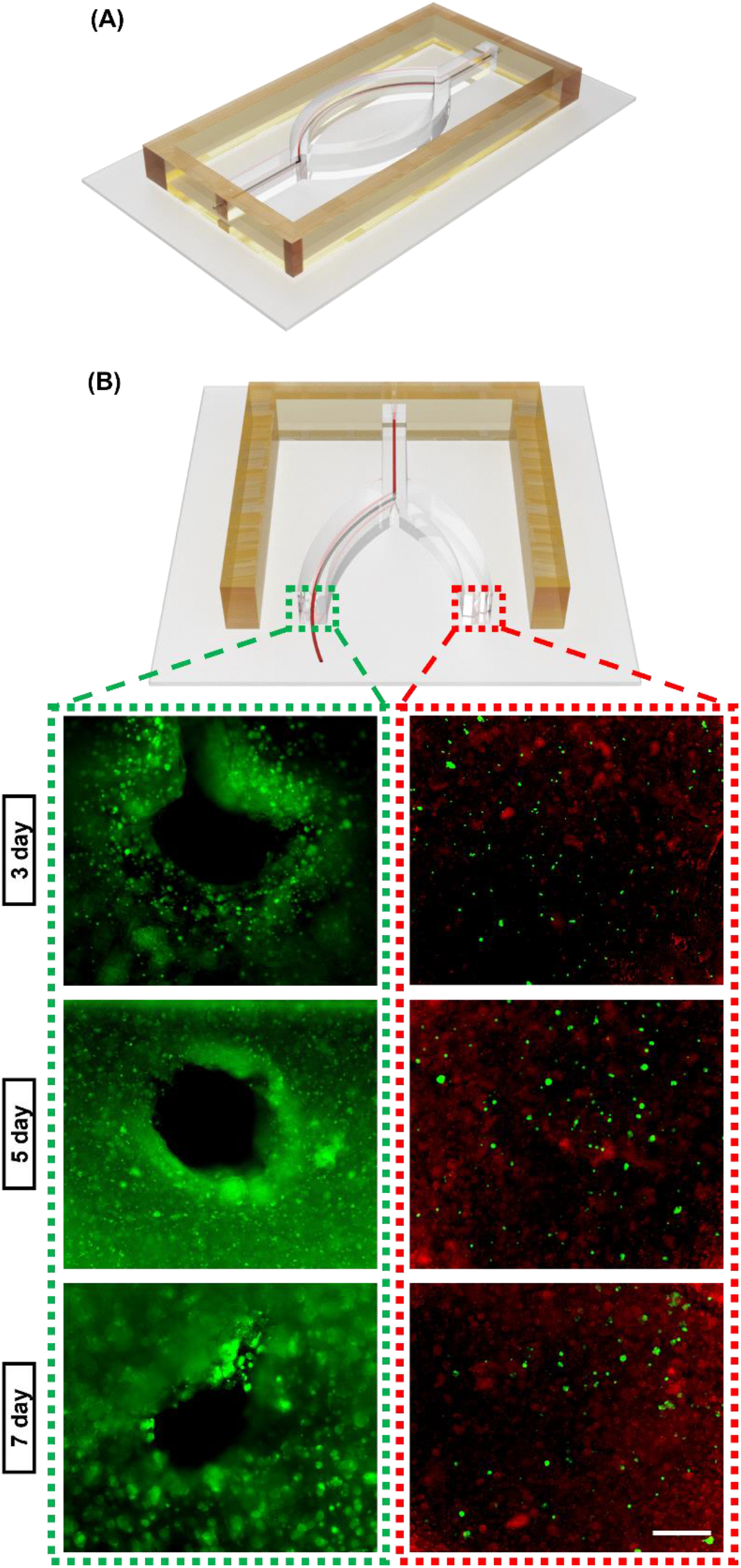
Bioprinting structure. (A) Bioprinted chip and (B) Live/dead assay of the fabricated chip to compare the perfused and unperfumed tissue models (scale: 250um)

## 4. Discussion

The idea of employing 3D bioprinting within supporting gel has garnered attention and discussion within the scientific community over the past several years. This approach has paved the way for the biofabrication of complex tissue and organ constructs, incorporating viable living cells. To effectively secure the locally extruded bioinks during the printing procedure, researchers have explored the utilization of diverse biocompatible pre-gels or hydrogel slurries exhibiting self-healing properties. Notably, substances such as hyaluronic acid [60] and gelatin [61,62] have been identified and employed to facilitate and enhance the bioprinting process. Nonetheless, when considering the utilization of the supporting bath as a permanent matrix, it becomes imperative to use a suitable method for crosslinking the bath material. In pursuit of this objective, photocrosslinking emerges as an enabling technique, affording the ability to fabricate more complex geometries. Within the scope of this investigation, GelMA-based hydrogels with nanoclay additives were introduced as a prospective matrix for the 3D printing of vascular structures. Recently, various studies have been utilized GelMA-LPN nanocomposites and investigated their characteristics and printability. Waters et al. [63] reported GelMA-LPN nanocomposite as growth factor carrier and controlled release vehicle. In another study, similar system was used as osteogenic and angiogenic scaffold [64]. In both of these studies, by the addition of LPN to GelMA, same trend of mechanical properties was reported as shown herein. The present findings clearly demonstrate that the inclusion of LPN exerts an inhibitory effect on the crosslinking process of the GelMA macromolecule [64] resulting in compression modulus decrease by adding 0.5% LPN to GelMA (Figure 1(B)). On the other hand, multitude of prior studies have consistently reported that the introduction of LPN leads to a substantial reinforcement of matrices [65–67]. As the concentration of LPN is incrementally increased, a notable trend emerges in the compression modulus, as illustrated in Figure 1(B). Specifically, in samples denoted as 5G1.5L and 5G2L, the compression modulus falls within the range exhibited by the 5G sample. This observation underscores that with the judicious addition of LPN to modulate the rheological characteristics, the resulting compression modulus closely approximates that of the 5G sample, indicating that the matrix permits vascular ingrowth to a sufficient degree [64]. Moreover, this mechanical behavior aligns with the microstructural evaluation. Notably, the 5G0.5L sample exhibits the largest pore size [68] (Figure 1(D) and **(E)**), which correlates with the lowest compression modulus and structural stability, thereby establishing a consistent relationship between microstructure and mechanical properties. Larger pore size also resulted in more water uptake capacity, thus 5G0.5L sample showed highest swelling ration and by enhancing LPN concentration, the swelling decreased as well. The influence of pore size on water uptake capacity is evident (Figure 1(C)), with larger pores, as exemplified by the 5G0.5L sample, displaying a heightened swelling ratio. Conversely, with the escalation of LPN concentration, a reduction in swelling becomes apparent. The extent of swelling also exerts a notable influence on the degradation process. In particular, the 5G0.5L sample, characterized by its highest swelling ratio and pore size, exhibits an increased water uptake within its 3D structure. Consequently, during the degradation study, this sample absorbs a greater quantity of water containing collagenase enzyme, leading to the highest degradation rate observed among the samples studied. Conversely, the 5G1.5L and 5G2L samples, featuring smaller pore sizes and lower swelling ratios, demonstrate reduced rates of degradation as shown in **Figure S2** [69]. This property renders them well-suited for more prolonged in vitro investigations, such as on-chip applications. It is worth noting that despite the similarity in pore size and swelling ratio between these two samples and the 5G control, they exhibit sustained weight retention after approximately 10 hours, in contrast to the 5G sample, which undergoes complete degradation within 5 hours. This phenomenon may be attributed to their structural compactness and the presence of LPN, which display greater resistance to degradation in a collagenase solution compared to GelMA (**Figure S2**).

The suitability of the developed nanocomposite hydrogel as a support matrix for hollow channel bioprinting was assessed, with a focus on key rheological properties, including shear-thinning, rapid gelling, and recovery behaviors. To understand the linear viscoelastic (LVE) range and yield point of the nanocomposite hydrogels, dynamic strain sweeps were conducted, revealing that all samples, except for 5G0L and 5G0.5L, demonstrated a wide range of LVE behavior as shown in Figure 2(A). As the concentration of LPN increased, the range of change in G’ and G” prior to the yield point decreased, suggesting that hydrogels containing more nanoclay exhibited increased elasticity and stability under oscillating strain. The onset of the decrease in G’ and increase in G” indicated damage to the inner structure of the hydrogels. Furthermore, all groups, except for 5G0L and 5G0.5L, exhibited clear yielding points at room temperature. Notably, the yielding strain decreased with increasing LPN concentration.

To understand the frequency dependence of the nanocomposite hydrogels, frequency sweep tests were performed within the LVE region. With the exception of the 5G0L and 5G0.5L samples, which had indistinguishable G’ and G”, other samples displayed G’ > G’’, indicating a predominantly elastic nature with minimal dependence on the applied frequency (Figure 2(B)). As the LPN concentration increased, the interactions between GelMA and LPN transitioned from a viscoelastic-dominant gel state to a glassy state colloidal network, where the complex modulus was nearly independent of the deformation frequency and showing extended plateau region. Also, increasing trend in G’ and G” is indicating more structural density regarding more LPN particles present in the matrix.

The viscosity of the inks was also determined, with samples featuring LPN concentrations above 0.5% exhibiting significantly higher viscosity compared to the 5G0L and 5G0.5L samples as illustrated in Figure 2(C) and by increasing the LPN concentration, the viscosity increases as well. The observed direct correlation between the increased viscosity and the concentration of LPN may be attributed to physical crosslinking. This phenomenon occurs as a result of interactions between the predominantly negatively charged LPN disks and the positive charges associated with the lysine ε-amino groups of GelMA [70]. Also, for these combinations, a shear thickening behavior is observed at lower shear rates which could be attributed to the unsteady state flow causing the experimental artifact [71]. As another aspect, these samples displayed shear-thinning behavior as the shear rate increased. Shear-thinning behavior is a crucial feature for extrusion 3D bioprinting, as well as for inks used in support structures for embedded printing. This behavior underscores the dynamic interactions between the polymer and Laponite, which can be dynamically formed and disrupted [72,73]. Thus, the shear-thinning property of GelMA-LPN nanocomposite hydrogel facilitates the passage of the needle through the supporting bath, which forms a stable hydrogel matrix due to its high viscosity when the shear stress is released.

Furthermore, it was observed that GelMA-LPN demonstrated structural recovery behavior after the removal of applied higher shear rates (Figure 2(D)). The thixotropic characteristics of the material allow the disrupted matrix to rebuild interactions over time, leading to the reformation of the matrix network. [74,75] This self-recovery property is an essential feature of the composite matrix for use as a support-bath material. Dynamic strain tests were conducted at both high (500%) and low (10%) strains. As shown in the graph, even within a short duration of 10 seconds, the structures with 1% and 1.5% of LPN demonstrated a significant recovery in storage modulus, almost returning to their initial values. However, the composite with 2% of LPN exhibited a lower recovery in storage modulus compared to the starting point. This difference may be attributed to the high amount of LPN, which may hinder the rapid recovery of GelMA chains, preventing them from efficiently reattaching to the charged surfaces of Laponite nanodiscs [76]. Through all these rheological characterizations, it was decided that 5G1.5L samples is showing the best properties to be used as support bath biomaterial for further bioprinting evaluations. Moreover, the rheological changes were analyzed during UV crosslinking of GelMA-LPN composite hydrogel (**Figure S4**). All GelMA-LPN composite hydrogels exhibited a crosslinking reaction plateau within 5-10 seconds. The difference between the crosslinking times could be attributed to the light scattering of nanoparticles as the amount of LPN increased, which may contribute to the reduction in crosslinking time. The saturation limit of the storage modulus (G’) shown in **Figure S4(A)** is in correlation with the mechanical properties previously reported and discussed. Furthermore, the storage modulus (G’) of the fully crosslinked GelMA-LPN samples was evaluated as a function of the angular frequency (ω) to investigate the effect of LPN on the mechanical behavior within the non-destructive strain range of the crosslinked composite hydrogel (**Figure S4(C)**). All the samples exhibited an independent complex modulus at high frequencies, and the amplitude of G’ of the crosslinked samples followed the same trend as observed in the mechanical evaluation. The addition of 0.5% LPN to the GelMA solution led to a decrease in storage modulus of at least ten times. As the LPN concentration increased, it acted as reinforcement particles, and the storage modulus approached the initial G’ observed in the control sample.

Additionally, in the gel-like state, which is the desired form of the bath material for bioprinting applications, it can be challenging to uniformly dispense cells. Aqueous dispersions of certain nanoclays are known to undergo aging and transition from a liquid-like state to a gel-like structure over time [77]. This means that the rheological parameters of an LPN gel, such as storage modulus, yield stress, or viscosity, can significantly increase with aging time [78]. Thus, it becomes possible to dispense cells in the liquid-like structure, and after pouring it into the appropriate mold, it will transform into a gel-like matrix after a resting period for the next printing step. A strain sweep measurement conducted on 5G0L and 5G1.5L samples right after vortexing and after 5 minutes of rest showed no change in the 5G0L sample, as there is no additional component to form an internal structure (**Figure S5**). However, in the case of the 5G1.5L sample, LPN formed an isotropic gel, with LPN partially interacting with each other in a ‘house of cards’ assembly, as illustrated in the schematic (**Figure S5**) [79]. This resulted in the expansion of the linear viscoelastic (LVE) range and a lower yield point at which the shear strains led to a more viscous region in the support bath.

In assessing another pivotal attribute of the supporting bath, we conducted an evaluation of its transparency. The results revealed a reduction in transmittance values with increasing LPN concentration. This phenomenon was primarily attributed to the choice of PBS as the solvent for the nanocomposite hydrogel solutions. It became evident that the presence of certain salts in PBS hindered the complete exfoliation of nanoclay platelets, causing them to form aggregates known as tactoids [41]. The observed decline in transparency is inherently linked to the light-scattering properties exhibited by these nanoclay tactoids. This behavior aligns with similar trends observed in various other nanomaterials [80,81]. Furthermore, this light scattering effect is expected to exert an influence on the crosslinking behavior. In the case of thin samples, our results demonstrated exceptional resolution in the SLA printing process. However, for thicker samples, the aforementioned light scattering properties may lead to over-crosslinking to some extent.

In printing results, it’s observed that the diameter of the printed structures is larger than the diameter of the nozzle that have been used (Figure 7(B)). It’s a common observation with hydrogels that the extruded filament diameter tends to be larger than the nozzle diameter. This phenomenon is typically attributed to the “Barus effect” or die swelling behavior, as discussed in section 3.4. This behavior can be explained by the fact that, while passing through a micron-sized nozzle, the polymer experiences shear stress, causing the polymer chains to stretch. When the material exits the nozzle, the removal of shear stress and a drop in pressure lead to the relaxation of these polymeric chains, resulting in the observed behavior. This phenomenon has been widely documented, especially in injection molding, and it depends on various factors, including ink temperature, material properties, extrusion speed, extrusion pressure, and nozzle geometry [82–85]. Furthermore, an increase in extrusion pressure is associated with higher material velocity as it exits the nozzle. Consequently, a larger volume of material is extruded, exceeding the nozzle diameter.

Conversely, at lower extrusion pressures, the material velocity compared to the printing speed is lower, resulting in stretched or inconsistent printing, as noted by Khalil et. al [86]. It’s also noteworthy that when the bath material has lower viscosity than the bioink being deposited within it, there tends to be more spreading after printing. Conversely, if the viscosity of the bath material is equal to or higher than that of the bioink, there is less or no spreading of the bioink, which contributes to better shape fidelity [87].

Overall, the developed bath material shows better recovery behavior, as depicted in the rheology tests (Figure 2d) and supports the printed structures well. This has been demonstrated by printing various channel structures having different levels of complexity; for instance, we printed simple straight lines, wavy lines, and structures having junctions, and later, also, through the photocrosslinking process, different shapes such as the liver structure was photo-patterned over it (**please refer to supplementary videos 1-5**). Further, to demonstrate the 3D structure support capability of the developed GelMA LPN bath material, we performed embedded printing of a coil-shaped 3D structure (**please refer to supplementary video 11**).

Regarding our biological study, it is worth noting that GelMA-LPN nanocomposite hydrogel biocompatibility has been extensively investigated in various studies [88,89], confirming its suitability for use as a bioink. However, we conducted a specific inquiry into the fate of HepG2 cells by encapsulating them within a 3D matrix of the 5G1.5L sample, identified as the optimal choice for use as a supporting bath. Barros et. al [89] findings indicate that LPN plays a role in upregulating the production of tumor cell growth factors, as well as genes associated with tissue remodeling and cell differentiation. Thus, in our study, HepG2 cells exhibit a heightened propensity to form larger and more numerous cell clusters. During the initial two weeks of culture, the 5G0L sample displayed a higher number of clusters, although they were comparatively smaller in size. Subsequently, the 5G1.5L sample exhibited not only larger clusters but also a greater quantity of clusters, attributed to enhanced cell proliferation in the presence of LPN.

Finally, the bioprinted structures reveal noteworthy findings regarding the cell viability within the channeled section. According to existing literature, it is established that cells located within a proximity of approximately 200 μm from a nutrient source can access an adequate supply of nutrients [90]. Consequently, the cells situated near the printed channel exhibit a markedly higher viability. Moreover, after a 7-day interval, HepG2 cells exhibit a tendency to form clusters and demonstrate robust proliferation. In stark contrast, cells within the solid section, positioned beyond the critical diffusion distance, manifest a pronounced increase in mortality rates, primarily attributable to an insufficient availability of essential nutrients and oxygen. These outcomes underscore the imperative need for vascularization within in vitro tissue constructs.

## 5. Conclusion

In conclusion, we successfully modified the rheological properties of GelMA to create a suitable support bath for embedded bioprinting. Material characterizations revealed that the 5G1.5L combination shows significant promise for this purpose. We achieved successful embedded printing of PF-127 as a sacrificial material in various structures using support bath containing 1.5% LPN. Furthermore, HepG2 cells proliferated well when encapsulated in the bioink we mentioned, and the channels printed within the structures played a crucial role in maintaining cell viability at over 95%. These results demonstrate that the optimized support bath bioink can serve as scaffolds for vascularized tissue fabrication in a variety of biological applications.

## Acknowledgement

This work was supported by a Natural Sciences and Engineering Research Council of Canada (NSERC) Discovery Grant (RGPIN-2014-04010) and Canada Foundation for Innovation John R. Evans Leaders Opportunity Fund.

## References

1. Esdaille, C.J., Washington, K.S., and Laurencin, C.T. (2021) Regenerative engineering: a review of recent advances and future directions. 10.2217/rme-2021-0016, 16 (5), 495–512.

2. Langer, R., and Vacanti, J.P. (1993) Tissue Engineering. Science (1979), 260 (5110), 920–926.

3. Song, H.H.G., Rumma, R.T., Ozaki, C.K., Edelman, E.R., and Chen, C.S. (2018) Vascular Tissue Engineering: Progress, Challenges, and Clinical Promise. Cell Stem Cell, 22 (3), 340– 354.

4. Carmeliet, P., and Jain, R.K. (2000) Angiogenesis in cancer and other diseases. Nature 2000 407:6801, 407 (6801), 249–257.

5. Devillard, C.D., and Marquette, C.A. (2021) Vascular Tissue Engineering: Challenges and Requirements for an Ideal Large Scale Blood Vessel. Front Bioeng Biotechnol, 9, 721843.

6. Tomasina, C., Bodet, T., Mota, C., Moroni, L., and Camarero-Espinosa, S. (2019) Bioprinting Vasculature: Materials, Cells and Emergent Techniques. Materials 2019, Vol. 12, Page 2701, 12 (17), 2701.

7. Tabatabaei Rezaei, N., Kumar, H., Liu, H., Lee, S.S., Park, S.S., and Kim, K. (2023) Recent Advances in Organ-on-Chips Integrated with Bioprinting Technologies for Drug Screening. Adv Healthc Mater, 12 (20), 2203172.

8. Awad, N.K., Niu, H., Ali, U., Morsi, Y.S., and Lin, T. (2018) Electrospun Fibrous Scaffolds for Small-Diameter Blood Vessels: A Review. Membranes 2018, Vol. 8, Page 15, 8 (1), 15.

9. Seifu, D.G., Purnama, A., Mequanint, K., and Mantovani, D. (2013) Small-diameter vascular tissue engineering. Nature Reviews Cardiology 2013 10:7, 10 (7), 410–421.

10. Quint, C., Kondo, Y., Manson, R.J., Lawson, J.H., Dardik, A., and Niklason, L.E. (2011) Decellularized tissue-engineered blood vessel as an arterial conduit. Proc Natl Acad Sci U S A, 108 (22), 9214–9219.

11. Best, C., Strouse, R., Hor, K., Pepper, V., Tipton, A., Kelly, J., Shinoka, T., and Breuer, C. (2018) Toward a patient-specific tissue engineered vascular graft. J Tissue Eng, 9.

12. Wang, Z., Mithieux, S.M., and Weiss, A.S. (2019) Fabrication Techniques for Vascular and Vascularized Tissue Engineering. Adv Healthc Mater, 8 (19), 1900742.

13. Zhu, J., Wang, Y., Zhong, L., Pan, F., and Wang, J. (2021) Advances in tissue engineering of vasculature through three-dimensional bioprinting. Developmental Dynamics, 250 (12), 1717–1738.

14. Bertassoni, L.E., Cecconi, M., Manoharan, V., Nikkhah, M., Hjortnaes, J., Cristino, A.L., Barabaschi, G., Demarchi, D., Dokmeci, M.R., Yang, Y., and Khademhosseini, A. (2014) Hydrogel bioprinted microchannel networks for vascularization of tissue engineering constructs. Lab Chip, 14 (13), 2202–2211.

15. Lee, V.K., Kim, D.Y., Ngo, H., Lee, Y., Seo, L., Yoo, S.S., Vincent, P.A., and Dai, G. (2014) Creating perfused functional vascular channels using 3D bio-printing technology. Biomaterials, 35 (28), 8092–8102.

16. Gao, G., Park, J.Y., Kim, B.S., Jang, J., and Cho, D.W. (2018) Coaxial Cell Printing of Freestanding, Perfusable, and Functional In Vitro Vascular Models for Recapitulation of Native Vascular Endothelium Pathophysiology. Adv Healthc Mater, 7 (23), 1801102.

17. Kolesky, D.B., Homan, K.A., Skylar-Scott, M.A., and Lewis, J.A. (2016) Three-dimensional bioprinting of thick vascularized tissues. Proc Natl Acad Sci U S A, 113 (12), 3179–3184.

18. Gioffredi, E., Boffito, M., Calzone, S., Giannitelli, S.M., Rainer, A., Trombetta, M., Mozetic, P., and Chiono, V. (2016) Pluronic F127 Hydrogel Characterization and Biofabrication in Cellularized Constructs for Tissue Engineering Applications. Procedia CIRP, 49, 125–132.

19. Suntornnond, R., An, J., and Chua, C.K. (2017) Bioprinting of Thermoresponsive Hydrogels for Next Generation Tissue Engineering: A Review. Macromol Mater Eng, 302 (1), 1600266.

20. Zhu, J., Wang, Y., Zhong, L., Pan, F., and Wang, J. (2021) Advances in tissue engineering of vasculature through three-dimensional bioprinting. Developmental Dynamics, 250 (12), 1717–1738.

21. Parak, A., Pradeep, P., du Toit, L.C., Kumar, P., Choonara, Y.E., and Pillay, V. (2019) Functionalizing bioinks for 3D bioprinting applications. Drug Discov Today, 24 (1), 198–205.

22. Choi, G., and Cha, H.J. (2019) Recent advances in the development of nature-derived photocrosslinkable biomaterials for 3D printing in tissue engineering. Biomater Res, 23 (1), 1– 7.

23. Choi, G., and Cha, H.J. (2019) Recent advances in the development of nature-derived photocrosslinkable biomaterials for 3D printing in tissue engineering. Biomater Res, 23 (1), 1– 7.

24. Parak, A., Pradeep, P., du Toit, L.C., Kumar, P., Choonara, Y.E., and Pillay, V. (2019) Functionalizing bioinks for 3D bioprinting applications. Drug Discov Today, 24 (1), 198–205.

25. Zhang, W., Ye, W., and Yan, Y. (2022) Advances in Photocrosslinkable Materials for 3D Bioprinting. Adv Eng Mater, 24 (1), 2100663.

26. Kumar, H., Sakthivel, K., Mohamed, M.G.A., Boras, E., Shin, S.R., and Kim, K. (2021) Designing Gelatin Methacryloyl (GelMA)-Based Bioinks for Visible Light Stereolithographic 3D Biofabrication. Macromol Biosci, 21 (1), 2000317.

27. Yue, K., Trujillo-de Santiago, G., Alvarez, M.M., Tamayol, A., Annabi, N., and Khademhosseini, A. (2015) Synthesis, properties, and biomedical applications of gelatin methacryloyl (GelMA) hydrogels. Biomaterials, 73, 254–271.

28. Wu, Q., Song, K., Zhang, D., Ren, B., Sole-Gras, M., Huang, Y., and Yin, J. (2022) Embedded extrusion printing in yield-stress-fluid baths. Matter, 5 (11), 3775–3806.

29. Li, Q., Ma, L., Gao, Z., Yin, J., Liu, P., Yang, H., Shen, L., and Zhou, H. (2022) Regulable Supporting Baths for Embedded Printing of Soft Biomaterials with Variable Stiffness. ACS Appl Mater Interfaces, 14 (37), 41695–41711.

30. Gaharwar, A.K., Cross, L.M., Peak, C.W., Gold, K., Carrow, J.K., Brokesh, A., Abhay Singh K Gaharwar, K.A., Cross, L.M., Peak, C.W., Gold, K., Carrow, J.K., Brokesh, A., Singh Biomedical Engineering K., and Gaharwar, A.K. (2019) 2D Nanoclay for Biomedical Applications: Regenerative Medicine, Therapeutic Delivery, and Additive Manufacturing. Advanced Materials, 31 (23), 1900332.

31. Nadernezhad, A., Caliskan, O.S., Topuz, F., Afghah, F., Erman, B., and Koc, B. (2019) Nanocomposite Bioinks Based on Agarose and 2D Nanosilicates with Tunable Flow Properties and Bioactivity for 3D Bioprinting. ACS Appl Bio Mater, 2 (2), 796–806.

32. Ding, H., and Chang, R.C. (2018) Printability Study of Bioprinted Tubular Structures Using Liquid Hydrogel Precursors in a Support Bath. Applied Sciences 2018, Vol. 8, Page 403, 8 (3), 403.

33. Cai, F.-F., Heid, S., Boccaccini, A.R., Aldo Boccaccini, C.R., and Y W O R D S Ada-, K.E. (2021) Potential of Laponite® incorporated oxidized alginate–gelatin (ADA-GEL) composite hydrogels for extrusion-based 3D printing. J Biomed Mater Res B Appl Biomater, 109 (8), 1090–1104.

34. Munoz-Perez, E., Perez-Valle, A., Igartua, M., Santos-Vizcaino, E., and Hernandez, R.M. (2023) High resolution and fidelity 3D printing of Laponite and alginate ink hydrogels for tunable biomedical applications. Biomaterials Advances, 149, 213414.

35. Boucenna, I., Royon, L., Colinart, P., Guedeau-Boudeville, M.A., and Mourchid, A. (2010) Structure and thermorheology of concentrated pluronic copolymer micelles in the presence of laponite particles. Langmuir, 26 (18), 14430–14436.

36. Chang, C.W., Van Spreeuwel, A., Zhang, C., and Varghese, S. (2010) PEG/clay nanocomposite hydrogel: a mechanically robust tissue engineering scaffold. Soft Matter, 6 (20), 5157–5164.

37. Kumar, H., Sakthivel, K., Mohamed, M.G.A., Boras, E., Shin, S.R., and Kim, K. (2021) Designing Gelatin Methacryloyl (GelMA)-Based Bioinks for Visible Light Stereolithographic 3D Biofabrication. Macromol Biosci, 21 (1), 2000317.

38. Zhang, K., Liu, B., Yang, Y., -, al, Miao, S., Cui, H., Nowicki, M., Wang, Z., Abdulla, R., Parker, B., Samanipour, R., Ghosh, S., and Kim, K. (2015) A simple and high-resolution stereolithography-based 3D bioprinting system using visible light crosslinkable bioinks. Biofabrication, 7 (4), 045009.

39. Wang, Z., Kumar, H., Tian, Z., Jin, X., Holzman, J.F., Menard, F., and Kim, K. (2018) Visible Light Photoinitiation of Cell-Adhesive Gelatin Methacryloyl Hydrogels for Stereolithography 3D Bioprinting. ACS Appl Mater Interfaces, 10 (32), 26859–26869.

40. Dawson, J.I., and Oreffo, R.O.C. (2013) Clay: New Opportunities for Tissue Regeneration and Biomaterial Design. Advanced Materials, 25 (30), 4069–4086.

41. Sheikhi, A., Afewerki, S., Oklu, R., Gaharwar, A.K., and Khademhosseini, A. (2018) Effect of ionic strength on shear-thinning nanoclay–polymer composite hydrogels. Biomater Sci, 6 (8), 2073–2083.

42. Bippus, L., Jaber, M., and Lebeau, B. (2009) Laponite and hybrid surfactant/laponite particles processed as spheres by spray-drying. New Journal of Chemistry, 33 (5), 1116–1126.

43. Ruzicka, B., and Zaccarelli, E. (2011) A fresh look at the Laponite phase diagram. Soft Matter, 7 (4), 1268–1286.

44. Ruzicka, B., Zulian, L., and Ruocco, G. (2004) Routes to gelation in a clay suspension. Phys Rev Lett, 93 (25), 258301.

45. Li, C., Mu, C., Lin, W., and Ngai, T. (2015) Gelatin Effects on the Physicochemical and Hemocompatible Properties of Gelatin/PAAm/Laponite Nanocomposite Hydrogels. ACS Appl Mater Interfaces, 7 (33), 18732–18741.

46. Pawar, N., and Bohidar, H.B. (2010) Spinodal decomposition and phase separation kinetics in nanoclay–biopolymer solutions. J Polym Sci B Polym Phys, 48 (5), 555–565.

47. Li, X., Liu, A., Ye, R., Wang, Y., and Wang, W. (2015) Fabrication of gelatin–laponite composite films: Effect of the concentration of laponite on physical properties and the freshness of meat during storage. Food Hydrocoll, 44, 390–398.

48. Wala, J., Maji, D., and Das, S. (2017) Influence of physico-mechanical properties of elastomeric material for different cell growth. Biomedical Materials, 12 (6), 065002.

49. Scognamiglio, F., Cok, M., Piazza, F., Marsich, E., Pacor, S., Aarstad, O.A., Aachmann, F.L., and Donati, I. (2023) Hydrogels based on methylated-alginates as a platform to investigate the effect of material properties on cell activity. The role of material compliance. Carbohydr Polym, 311, 120745.

50. Peppas, N.A., Hilt, J.Z., Khademhosseini, A., and Langer, R. (2006) Hydrogels in Biology and Medicine: From Molecular Principles to Bionanotechnology. Advanced Materials, 18 (11), 1345–1360.

51. McCormack, A., Highley, C.B., Leslie, N.R., and Melchels, F.P.W. (2020) 3D Printing in Suspension Baths: Keeping the Promises of Bioprinting Afloat. Trends Biotechnol, 38 (6), 584–593.

52. Grosskopf, A.K., Truby, R.L., Kim, H., Perazzo, A., Lewis, J.A., and Stone, H.A. (2018) Viscoplastic Matrix Materials for Embedded 3D Printing. ACS Appl Mater Interfaces, 10 (27), 23353–23361.

53. Yang, K., Liu, Z., Wang, J., and Yu, W. (2018) Stress bifurcation in large amplitude oscillatory shear of yield stress fluids. J Rheol (N Y N Y), 62 (1), 89–106.

54. Dávila, J.L., and d’Ávila, M.A. (2017) Laponite as a rheology modifier of alginate solutions: Physical gelation and aging evolution. Carbohydr Polym, 157, 1–8.

55. Jin, Y., Compaan, A., Chai, W., and Huang, Y. (2017) Functional Nanoclay Suspension for Printing-Then-Solidification of Liquid Materials. ACS Appl Mater Interfaces, 9 (23), 20057– 20066.

56. Zhang, C., Zhou, Y., Han, H., Zheng, H., Xu, W., and Wang, Z. (2021) Dopamine-Triggered Hydrogels with High Transparency, Self-Adhesion, and Thermoresponse as Skinlike Sensors. ACS Nano, 15 (1), 1785–1794.

57. He, Y., Yu, R., Li, X., Zhang, M., Zhang, Y., Yang, X., Zhao, X., and Huang, W. (2021) Digital Light Processing 4D Printing of Transparent, Strong, Highly Conductive Hydrogels. ACS Appl Mater Interfaces, 13 (30), 36286–36294.

58. Shin, S., Kwak, H., and Hyun, J. (2018) Melanin Nanoparticle-Incorporated Silk Fibroin Hydrogels for the Enhancement of Printing Resolution in 3D-Projection Stereolithography of Poly(ethylene glycol)-Tetraacrylate Bio-ink. ACS Appl Mater Interfaces, 10 (28), 23573– 23582.

59. Paxton, N., Smolan, W., Böck, T., Melchels, F., Groll, J., and Jungst, T. (2017) Proposal to assess printability of bioinks for extrusion-based bioprinting and evaluation of rheological properties governing bioprintability. Biofabrication, 9 (4), 044107.

60. Hoon Song, K., Highley, C.B., Rouff, A., Burdick, J.A., Song, K.H., Highley, C.B., Rouff, A., and Burdick, J.A. (2018) Complex 3D-Printed Microchannels within Cell-Degradable Hydrogels. Adv Funct Mater, 28 (31), 1801331.

61. Lee, A., Hudson, A.R., Shiwarski, D.J., Tashman, J.W., Hinton, T.J., Yerneni, S., Bliley, J.M., Campbell, P.G., and Feinberg, A.W. (2019) 3D bioprinting of collagen to rebuild components of the human heart. Science (1979), 365 (6452), 482–487.

62. Choi, Y.J., Jun, Y.J., Kim, D.Y., Yi, H.G., Chae, S.H., Kang, J., Lee, J., Gao, G., Kong, J.S., Jang, J., Chung, W.K., Rhie, J.W., and Cho, D.W. (2019) A 3D cell printed muscle construct with tissue-derived bioink for the treatment of volumetric muscle loss. Biomaterials, 206, 160– 169.

63. Waters, R., Pacelli, S., Maloney, R., Medhi, I., Ahmed, R.P.H., and Paul, A. (2016) Stem cell secretome-rich nanoclay hydrogel: a dual action therapy for cardiovascular regeneration. Nanoscale, 8 (14), 7371–7376.

64. Cidonio, G., Alcala-Orozco, C.R., Lim, K.S., Glinka, M., Mutreja, I., Kim, Y.H., Dawson, J.I., Woodfield, T.B.F., and Oreffo, R.O.C. (2019) Osteogenic and angiogenic tissue formation in high fidelity nanocomposite Laponite-gelatin bioinks. Biofabrication, 11 (3), 035027.

65. Xavier, J.R., Thakur, T., Desai, P., Jaiswal, M.K., Sears, N., Cosgriff-Hernandez, E., Kaunas, R., and Gaharwar, A.K. (2015) Bioactive nanoengineered hydrogels for bone tissue engineering: A growth-factor-free approach. ACS Nano, 9 (3), 3109–3118.

66. Chimene, D., Peak, C.W., Gentry, J.L., Carrow, J.K., Cross, L.M., Mondragon, E., Cardoso, G.B., Kaunas, R., and Gaharwar, A.K. (2018) Nanoengineered Ionic-Covalent Entanglement (NICE) Bioinks for 3D Bioprinting. ACS Appl Mater Interfaces, 10 (12), 9957–9968.

67. Paul, A., Manoharan, V., Krafft, D., Assmann, A., Uquillas, J.A., Shin, S.R., Hasan, A., Hussain, M.A., Memic, A., Gaharwar, A.K., and Khademhosseini, A. (2016) Nanoengineered biomimetic hydrogels for guiding human stem cell osteogenesis in three dimensional microenvironments. J Mater Chem B, 4 (20), 3544–3554.

68. Chimene, D., Peak, C.W., Gentry, J.L., Carrow, J.K., Cross, L.M., Mondragon, E., Cardoso, G.B., Kaunas, R., and Gaharwar, A.K. (2018) Nanoengineered Ionic-Covalent Entanglement (NICE) Bioinks for 3D Bioprinting. ACS Appl Mater Interfaces, 10 (12), 9957–9968.

69. Barros, N.R. de, Gomez, A., Ermis, M., Falcone, N., Haghniaz, R., Young, P., Gao, Y., Aquino, A.-F., Li, S., Niu, S., Chen, R., Huang, S., Zhu, Y., Eliahoo, P., Sun, A., Khorsandi, D., Kim, J., Kelber, J., Khademhosseini, A., Kim, H.-J., and Li, B. (2023) Gelatin methacryloyl and Laponite bioink for 3D bioprinted organotypic tumor modeling. Biofabrication, 15 (4), 045005.

70. Waters, R., Alam, P., Pacelli, S., Chakravarti, A.R., Ahmed, R.P.H., and Paul, A. (2018) Stem cell-inspired secretome-rich injectable hydrogel to repair injured cardiac tissue. Acta Biomater, 69, 95–106.

71. Wagner, C.E., Barbati, A.C., Engmann, J., Burbidge, A.S., and McKinley, G.H. (2016) Apparent shear thickening at low shear rates in polymer solutions can be an artifact of non-equilibration. Applied Rheology, 26 (5), 36–40.

72. Wilson, S.A., Cross, L.M., Peak, C.W., and Gaharwar, A.K. (2017) Shear-Thinning and Thermo-Reversible Nanoengineered Inks for 3D Bioprinting. ACS Appl Mater Interfaces, 9 (50), 43449–43458.

73. Peak, C.W., Stein, J., Gold, K.A., and Gaharwar, A.K. (2018) Nanoengineered Colloidal Inks for 3D Bioprinting. Langmuir, 34 (3), 917–925.

74. Dávila, J.L., and d’Ávila, M.A. (2017) Laponite as a rheology modifier of alginate solutions: Physical gelation and aging evolution. Carbohydr Polym, 157, 1–8.

75. Jin, Y., Compaan, A., Chai, W., and Huang, Y. (2017) Functional Nanoclay Suspension for Printing-Then-Solidification of Liquid Materials. ACS Appl Mater Interfaces, 9 (23), 20057– 20066.

76. Yang, K., Liu, Z., Wang, J., and Yu, W. (2018) Stress bifurcation in large amplitude oscillatory shear of yield stress fluids. J Rheol (N Y N Y), 62 (1), 89–106.

77. Shahin, A., and Joshi, Y.M. (2010) Irreversible aging dynamics and generic phase behavior of aqueous suspensions of laponite. Langmuir, 26 (6), 4219–4225.

78. Au, P.I., Hassan, S., Liu, J., and Leong, Y.K. (2015) Behaviour of LAPONITE® gels: Rheology, ageing, pH effect and phase state in the presence of dispersant. Chemical Engineering Research and Design, 101, 65–73.

79. Ruzicka, B., Zaccarelli, E., Zulian, L., Angelini, R., Sztucki, M., Moussaïd, A., Narayanan, T., and Sciortino, F. (2010) Observation of empty liquids and equilibrium gels in a colloidal clay. Nature Materials 2011 10:1, 10 (1), 56–60.

80. Ye, Y., Zhang, Y., Chen, Y., Han, X., Jiang, F., Ye, Y., Zhang, Y., Chen, Y., Han, X., and Jiang, F. (2020) Cellulose Nanofibrils Enhanced, Strong, Stretchable, Freezing-Tolerant Ionic Conductive Organohydrogel for Multi-Functional Sensors. Adv Funct Mater, 30 (35), 2003430.

81. Kim, S.W., and Cha, S.H. (2014) Thermal, mechanical, and gas barrier properties of ethylene– vinyl alcohol copolymer-based nanocomposites for food packaging films: Effects of nanoclay loading. J Appl Polym Sci, 131 (11), 40289.

82. Lee, J.M., and Yeong, W.Y. (2015) A preliminary model of time-pressure dispensing system for bioprinting based on printing and material parameters. Virtual Phys Prototyp, 10 (1), 3–8.

83. Cooke, M.E., and Rosenzweig, D.H. (2021) The rheology of direct and suspended extrusion bioprinting. APL Bioeng, 5 (1), 11502.

84. Vlachopoulos, J., and Polychronopoulos, N. (2011) Basic Concepts in Polymer Melt Rheology and Their Importance in Processing. Applied Polymer Rheology: Polymeric Fluids with Industrial Applications, 1–27.

85. Wu, D., Yu, Y., Tan, J., Huang, L., Luo, B., Lu, L., and Zhou, C. (2018) 3D bioprinting of gellan gum and poly (ethylene glycol) diacrylate based hydrogels to produce human-scale constructs with high-fidelity. Mater Des, 160, 486–495.

86. Khalil, S., and Sun, W. (2007) Biopolymer deposition for freeform fabrication of hydrogel tissue constructs. Materials Science and Engineering: C, 27 (3), 469–478.

87. Jin, Y., Chai, W., and Huang, Y. (2017) Printability study of hydrogel solution extrusion in nanoclay yield-stress bath during printing-then-gelation biofabrication. Materials Science and Engineering: C, 80, 313–325.

88. Dong, L., Bu, Z., Xiong, Y., Zhang, H., Fang, J., Hu, H., Liu, Z., and Li, X. (2021) Facile extrusion 3D printing of gelatine methacrylate/Laponite nanocomposite hydrogel with high concentration nanoclay for bone tissue regeneration. Int J Biol Macromol, 188, 72–81.

89. de Barros, N.R., Gomez, A., Ermis, M., Falcone, N., Haghniaz, R., Young, P., Gao, Y., Aquino, A.F., Li, S., Niu, S., Chen, R.R., Huang, S., Zhu, Y., Eliahoo, P., Sun, A., Khorsandi, D., Kim, J., Kelber, J., Khademhosseini, A., Kim, H.J., and Li, B. (2023) Gelatin methacryloyl and Laponite bioink for 3D bioprinted organotypic tumor modeling. Biofabrication, 15 (4), 045005.

90. Rouwkema, J., FJM Koopman, B., Van Blitterswijk, C.A., Dhert, W.J., and Malda, J. (2009) Supply of Nutrients to Cells in Engineered Tissues. Biotechnol Genet Eng Rev, 26 (1), 163– 178.

