## Supplementary file for "Vascularized Liver Tissue Embedded Bioprinting Utilizing GelMA/Nanoclay-based Composite hydrogels"

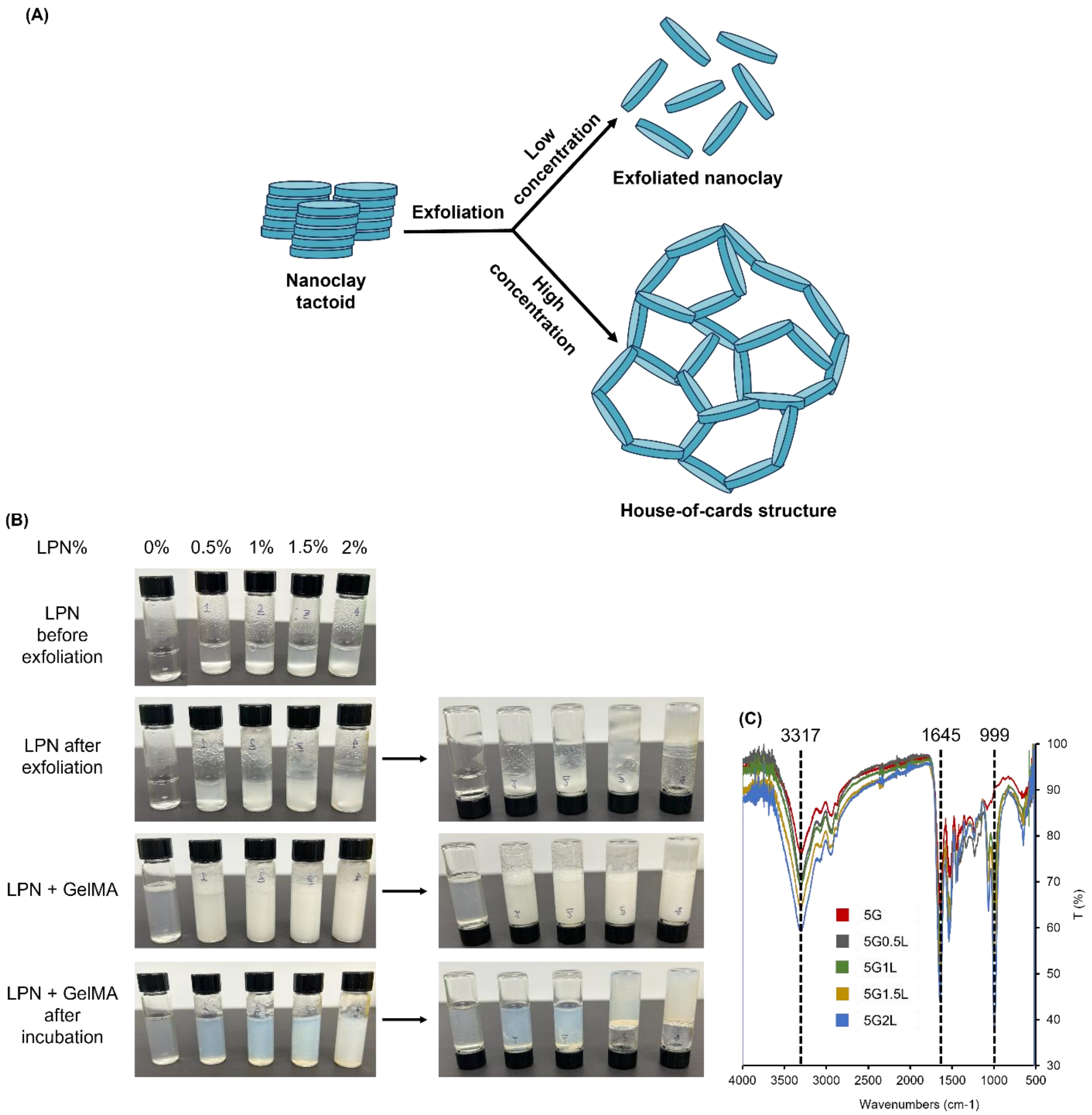


**Figure S 1** (A) schematic of the LPN particles distribution in solution: at low and high concentration solutions, (B) pictures showing the process of the of preparing GelMA-LPN nanocomposite hydrogels: i) LPN solution before exfoliation, ii) LPN solution after exfoliation, iii)LPN solution after adding GELMA solution, and iv) GelMA-LPN hydrogels after incubation. (C) FTIR spectrum of GelMA-LPN hydrogels

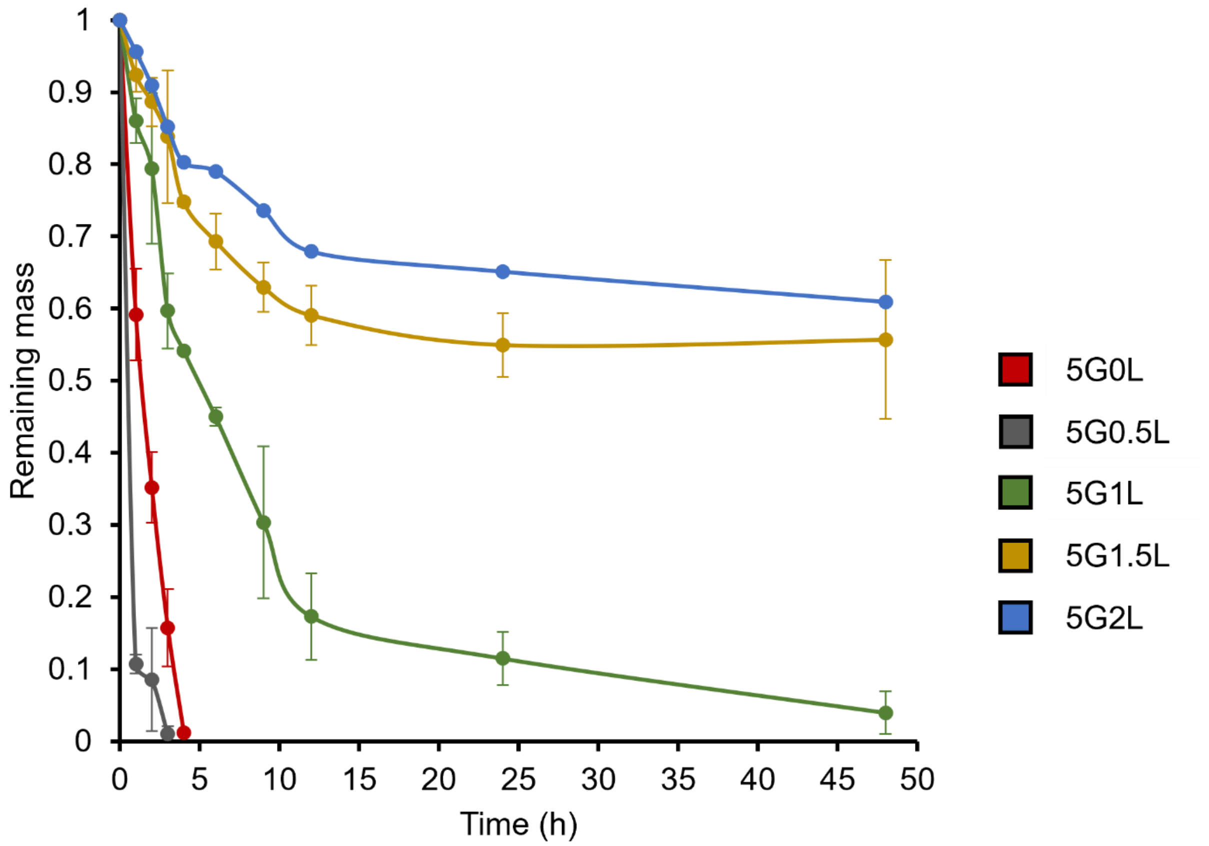


**Figure S 2** Degradation study of crosslinked GelMA-LPN nanocomposite hydrogels under enzymatic degradation condition

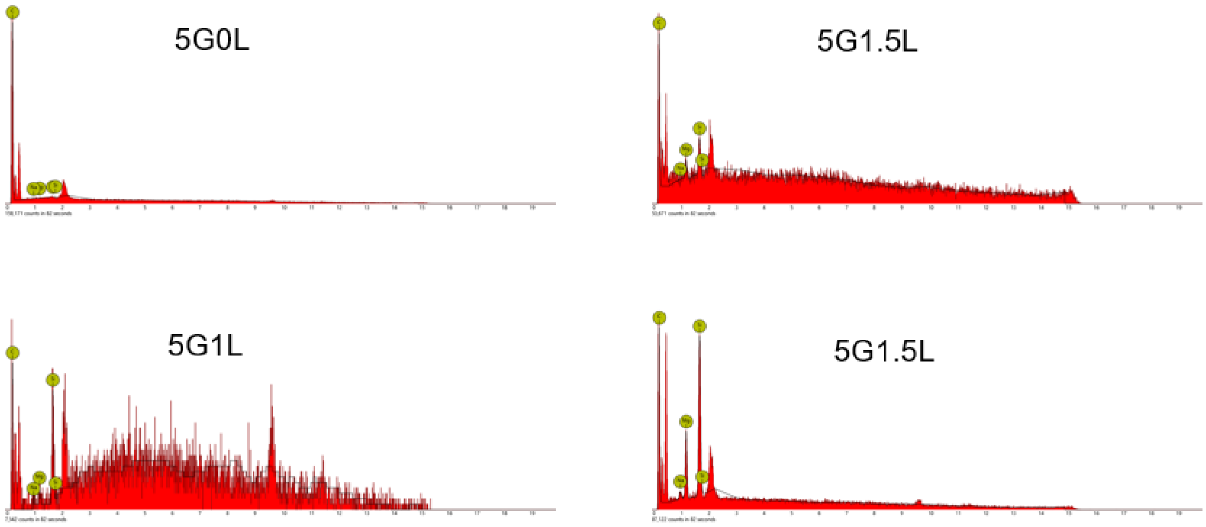

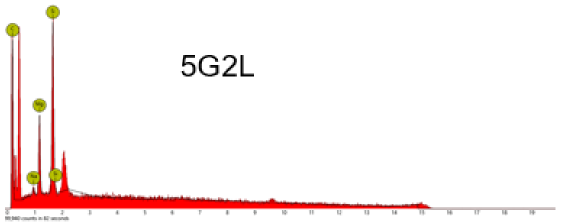


| LPN% | Si% |
|---|---|
| 0% | 0% |
| 0.5% | 2.29% |
| 1% | 5.05% |
| 1.5% | 5.38% |
| 2% | 5.87% |

**Figure S 3** EDX analysis to evaluate the presence of LPN in the GelMA-LPN hydrogel samples

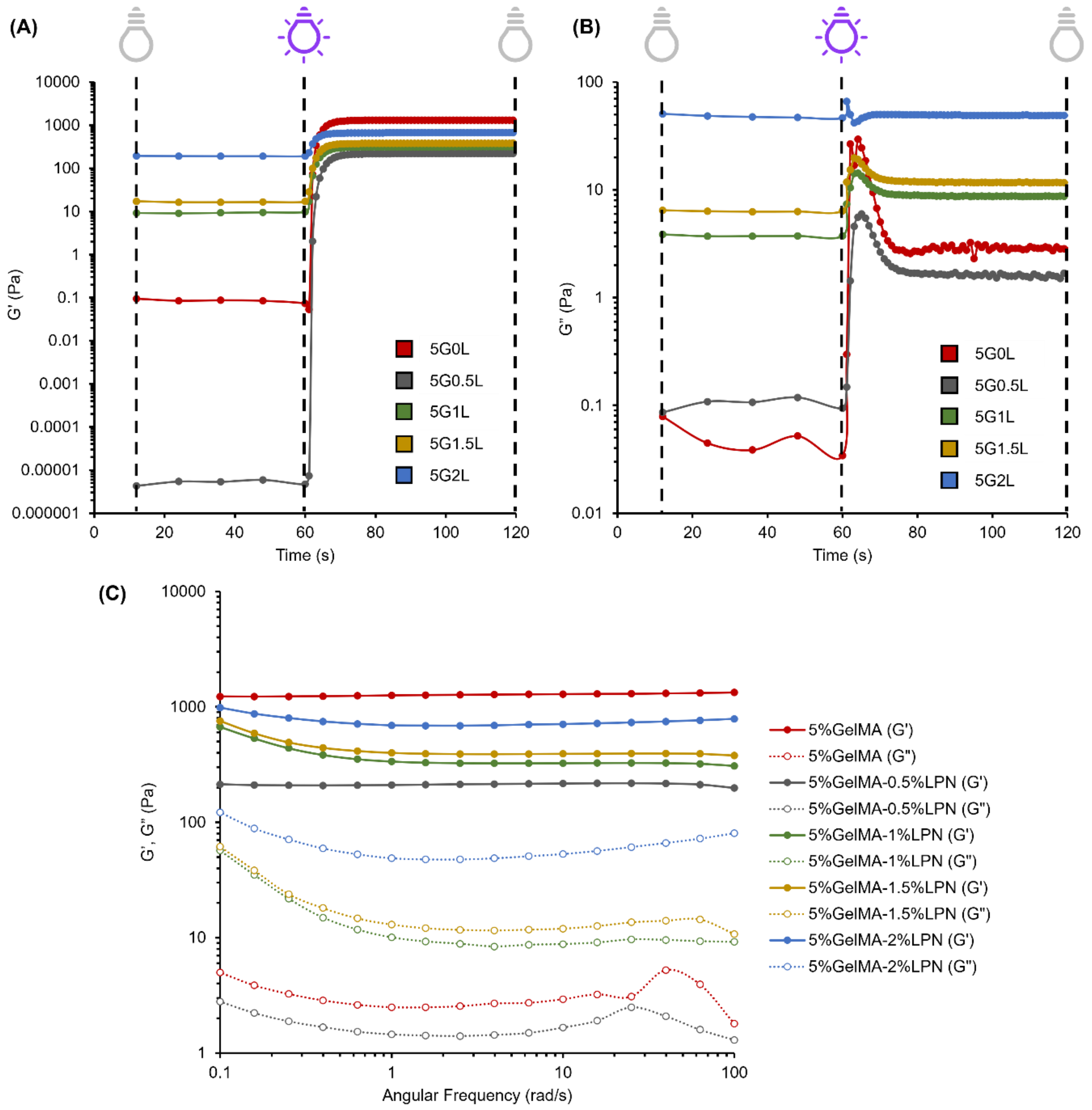


**Figure S 4** Kinetics of photocrosslinking evaluation of GelMA-LPN hydrogels. (A) storage modulus change by irradiation time, (B) loss modulus change by irradiation time, (C) Frequency sweep profiles of photocrosslinked inks

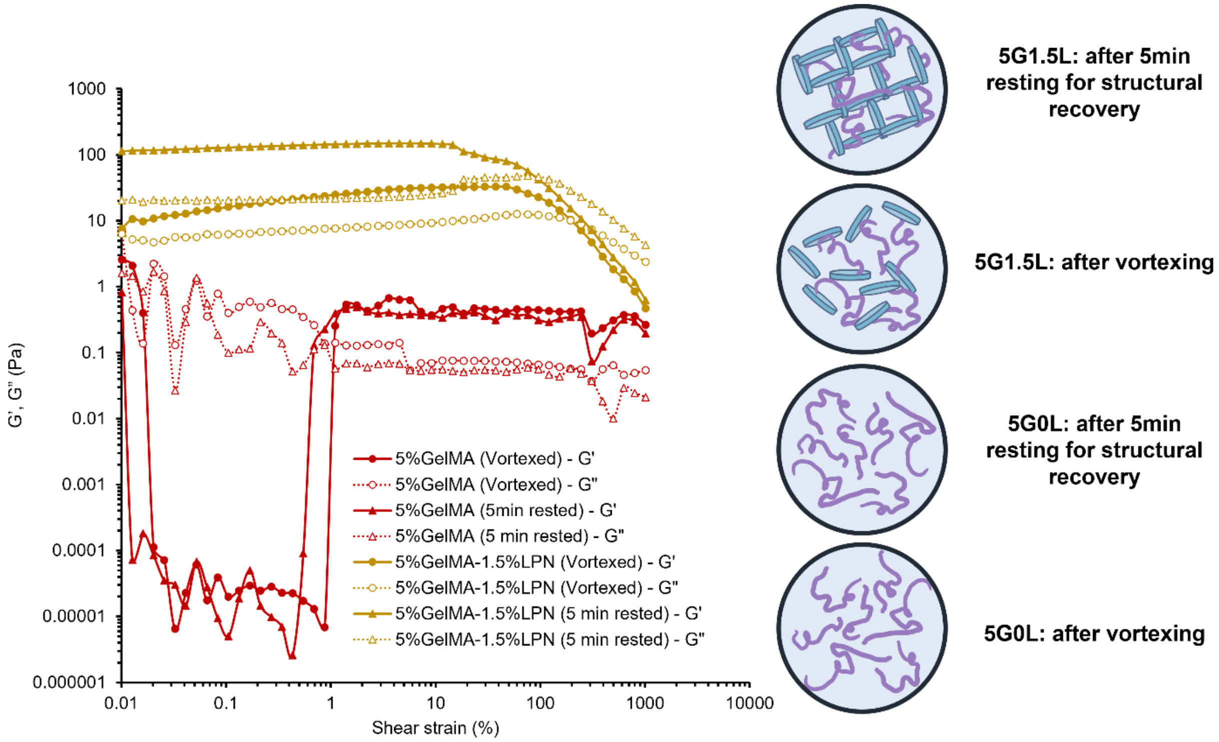


**Figure S 5** Strain amplitude sweep profiles of GelMA-LPN hydrogels to investigate the time dependant rheological properties

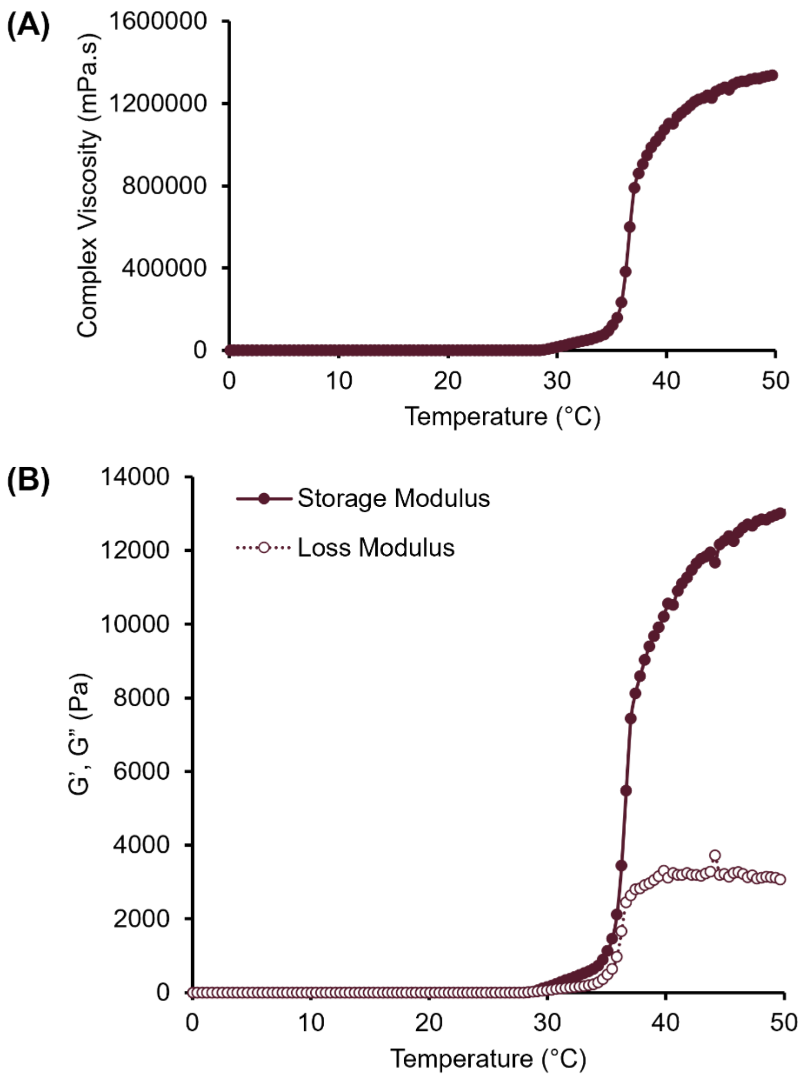


**Figure S 6** (A) Complex viscosity, (B) storage and loss modulus change of PF127 with temperature

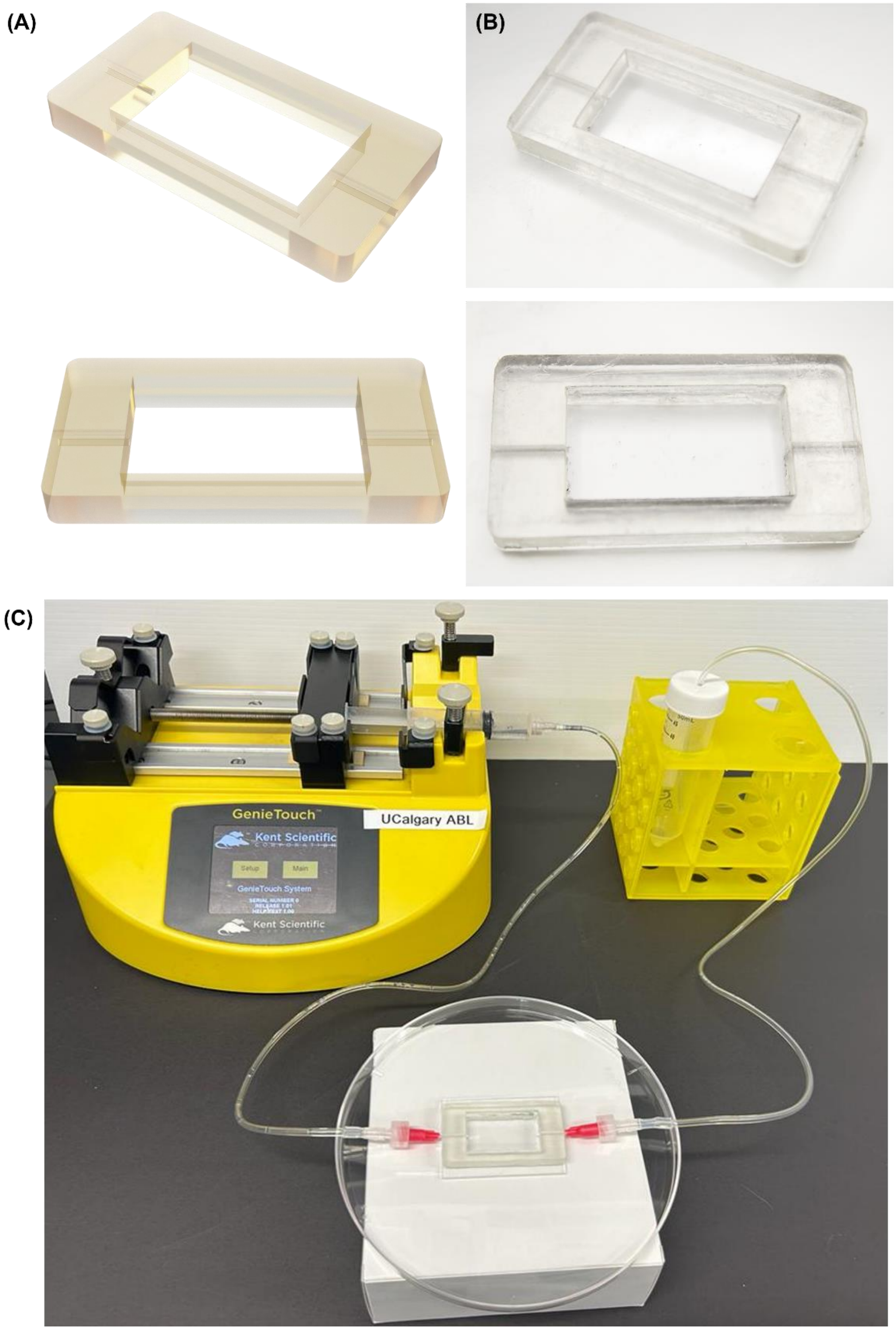

**Figure S 7** Molds used for the chip: (A) schematics and (B) real pics, (C) the setup for perfusion

**Table 1** Supplementary videos

| No. | Video title |
|---|---|
| 1 | Supplementary video 1 - Structure 1 embedded printing |
| 2 | Supplementary video 2 - Structure 2 embedded printing |
| 3 | Supplementary video 3 - Structure 3 embedded printing |
| 4 | Supplementary video 4 - Structure 4 embedded printing |
| 5 | Supplementary video 5 - Structure 5 embedded printing |
| 6 | Supplementary video 6 - 25PSI printing optimization |
| 7 | Supplementary video 7 - 30PSI printing optimization |
| 8 | Supplementary video 8 - 40PSI printing optimization |
| 9 | Supplementary video 9 - 50PSI printing optimization |
| 10 | Supplementary video 10 - 60PSI printing optimization |
| 11 | Supplementary video 11 - 3D coil printing |
